# Somatic variant detection using an individual-specific pangenome reference: a case study of a pediatric acute myeloid leukemia patient

**DOI:** 10.64898/2026.10.27.756267

**Authors:** Elizaveta Kulaeva, Eline JM Bertrums, Mark van Roosmalen, Rico Hagelaar, Lucca LM Derks, Laurianne Trabut, Arianne M Brandsma, Mark Verheul, Matteo Boretto, Sjors Middelkamp, Pjotr Prins, Ruben van Boxtel, Andrea Guarracino

## Abstract

**Background:** Reference bias can cause false or missed somatic variant calls, also in clinically relevant genes. Pangenomes aim to reduce this bias, but evidence of their diagnostic benefit is lacking. We tested how an individual-specific pangenome affects somatic variant calling in bulk samples and single cells.

**Results:** We built an individual-specific pangenome from Nanopore assemblies of normal hematopoietic stem and progenitor cells of a child with acute myeloid leukemia. We aligned short-read whole-genome sequencing data of bulk and single-cell samples to hg38 and to the pangenome, and called somatic variants with standard downstream tools after anchoring pangenome alignments to hg38. Single nucleotide variants were mostly shared, while indel calls unique to the pangenome were fewer than those unique to hg38, mostly in repeats. Both approaches detected *NUP98-NSD1* and *FLT3* internal tandem duplication (*FLT3*-ITD) drivers. The pangenome-based phylogeny showed more parallel branching in comparison to hg38-based tree. The pangenome removed three false-positive *PRSS1* variants and, in single cells, frameshift insertion calls next to polyG and polyC repeats. However, surjection to hg38 coordinates split a *WT1* driver indel call in a low-complexity region into two separate calls. Validation showed that all somatic structural variant calls, except *NUP98-NSD1* and *FLT3*-ITD, were germline variants or artifacts.

**Conclusions:** The use of individual-specific pangenome led to fewer false-positive indel and driver calls, mainly in repeats and single cells, and changed the branching of the inferred phylogeny. However, surjection artifacts and false-positive somatic structural variants show that pangenome-based results need read-level inspection and validation before clinical use.

## Background

Whole genome sequencing (WGS) is frequently used in diagnosis of cancer and rare diseases [1]. A fundamental limitation of this method lies in the reference genome concept itself. Most WGS analyses align reads against a single linear reference, such as hg38, and approximately 70% of this reference comes from one donor [2,3]. Reads from the regions where an individual’s genome diverges from the reference are harder to map correctly. In such cases, the aligner may misplace these reads, or fail to map them at all, creating reference bias [4] and unreliable variant calls, especially for individuals whose genomes are poorly represented in hg38 [3]. This problem becomes even more serious in somatic variant calling. If complex germline variants are missed due to poor alignment against the reference, they can be mistaken for somatic events, or true somatic mutations in these regions can be undetected [5,6]. Together, these limitations reduce both the sensitivity and the precision of somatic variant detection.

One strategy to mitigate the reference bias is to incorporate the missing variation into the reference genome using pangenomic approaches. Pangenomes represent the variation in a population as sequence graphs and better preserve intra-species genomic diversity in comparison to a single linear reference [7,8]. Graph-based aligners that map reads against a pangenome reference are more accurate than linear aligners at capturing germline variation across individuals [9]. However, a large pangenome graph containing hundreds of haplotypes is complex, and its variation absent from the sequenced genome can cause misalignment of that individual’s reads [10]. A more targeted alternative is an individual-specific pangenome, constructed from an individual’s own genome assembly anchored to a standard reference. Haplotype sampling, which keeps only the population haplotypes that match a sample’s reads, follows a similar idea [9].

Previous studies have already shown that pangenome references improve the identification of complex structural variants (SVs) that are often missed by traditional linear references. For instance, using a graph-based pangenome from 185 gastric cancer patients increased the total number of somatic SV calls from 21,415 with hg38 to 35,227 and found five candidate driver genes not detected with hg38, but missed nine detected with it [11]. Similarly, in autism research, pangenome assemblies used as controls filtered out over 97% of common SVs, enabling the prioritization of rare, potentially pathogenic mutations in previously unsolved cases [12].

At the somatic level, the human pangenome reference improved somatic single nucleotide variant (SNV) detection in 30 whole-exome-sequenced bladder tumors, with the largest gain in individuals of East Asian ancestry, by reducing germline contamination and reference bias [3]. Diploid donor-specific assemblies used as references found over 20% more somatic SVs than hg38 and T2T-CHM13 in six tumor-normal cell line pairs [5], and combining pangenome alignment with a de novo germline assembly reduced false-positive somatic SVs in long-read data [6]. In the melanoma cell line COLO829 and in a HapMap mixture dataset, a pangenome-inferred donor-specific assembly improved read mappability and somatic variant detection compared with hg38 [10]. Aligning to this assembly found 4,290 somatic SNVs not called with hg38, against 930 called only with hg38, and the assembly-only calls carried a higher proportion of UV light-associated single base substitution (SBS) signatures.

Despite the growing body of pangenomic research, personalized references have so far been tested mostly on cell lines, cell-line mixtures or healthy tissues and on three pediatric B-ALL samples with personalized diploid assemblies, while other patient tumors have been analyzed with population-based pangenomes [3,10,11,13]. To our knowledge, a pangenome built from a patient’s own assembly has not yet been tested for somatic short variants and SVs in primary cancer samples and single cells. We take a first step toward addressing this gap by incorporating a haplotype-resolved individual-specific pangenome reference into a somatic variant detection framework for a pediatric acute myeloid leukemia (pAML) case. pAML is an aggressive malignancy that accounts for 15 to 20% of all pediatric leukemia diagnoses in children under the age of 15 years [14]. Compared with adult AML, pAML is more often driven by SVs and chromosomal rearrangements, which were found in 70.7% of pAML cases [15,16]. This landscape, however, varies across populations [17], which limits how well any single reference or fixed driver catalog represents an individual patient.

Here we compare this pangenome reference with hg38 for the detection of somatic SNVs, insertion-deletion variants (indels) and SVs, by genomic context and driver impact. We validate selected calls by PCR and Sanger sequencing. This work assesses the practical value of individual-specific pangenome references in a clinical pediatric cancer setting, from variant detection to single-cell phylogeny reconstruction.

## Results

### Individual-specific pangenome construction and read alignment: from linear mappings through graph and back

Clinical diagnostic material used for this study was obtained from a female patient with primary pediatric cytogenetically normal AML and a bone marrow blast content of 77%. The blasts harbored two diagnostic driver events: a *NUP98-NSD1* fusion (confirmed by retrospective bulk RNA-seq analysis) and *FLT3*-ITD (confirmed by PCR), a combination associated with poor outcome [18]. To enable individual-specific pangenome construction while avoiding leukemic contamination, hematopoietic stem and progenitor cells (HSPCs) were isolated by fluorescence-activated cell sorting (FACS) using blast-negative gating. Sorted HSPCs were then expanded in bulk in a culture medium that suppresses blast proliferation (see Methods: *FACS and HSPC culture* section and [19]). We sequenced these HSPCs with Oxford Nanopore Simplex at 30x coverage, chosen as a practical compromise between cost and assembly quality in a clinical setting, and assembled the patient’s genome from these reads; along with that we sequenced FACS-sorted AML blasts at 15x coverage (Fig.1A, panel i).

**Figure 1:**
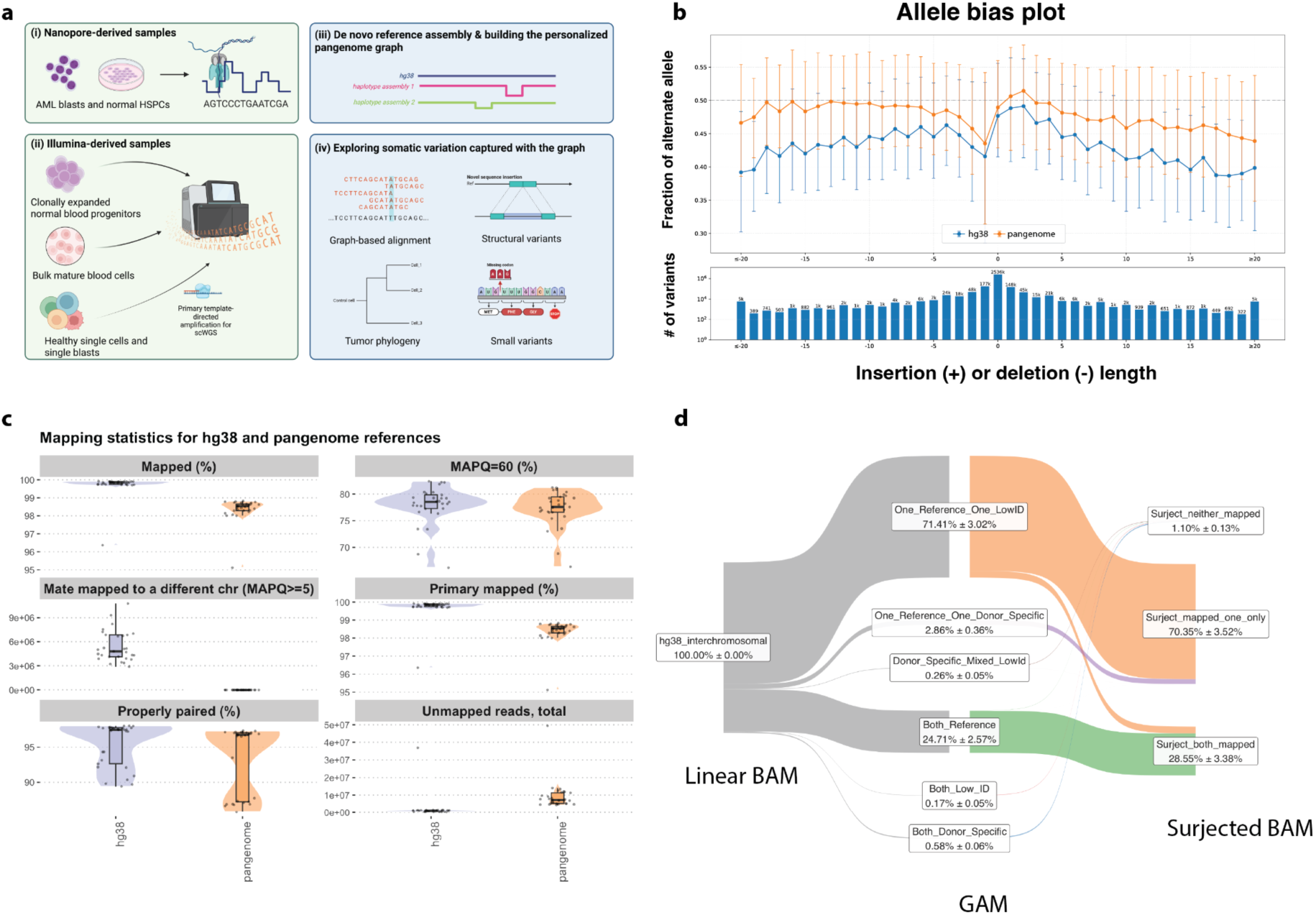
Evaluation of the pangenome assembly and of alignments to hg38 and to the pangenome references. **(A)** i. Long-read Nanopore sequencing (30x coverage) of expanded patient-derived normal hematopoietic stem and progenitor cells (HSPCs) and sorted AML blasts (15x coverage). ii Short-read Illumina sequencing of normal HSPCs and multipotent progenitors (clonally expanded) and single-cell populations (differentiated progenitors and tumor blasts) processed using PTA. iii. Construction of the individual-specific pangenome graph using Minigraph-Cactus, integrating hg38 with patient-specific *de novo* haplotype assemblies. iv. Graph-based alignment of short-read data to identify somatic small variants and SVs, allowing high-resolution tumor phylogenetic reconstruction. **(B)** Bias-by-allele-length plot of hg38 (blue) and pangenome (orange) alignments for the AML bulk sample. **(C)** Mapping quality metrics for linear and graph-based alignments for all samples. Violins are colored by the genomic reference used for mapping. **(D)** Pangenome tracking network of split high- confidence reads for bulk samples. A Sankey diagram illustrates the proportions of reads defined as reference-mapped, donor-specific, or low-identity within the graph, alongside their mapping labels after surjection. Values indicate the cohort mean percentage ± standard deviation across five bulk samples.

To comprehensively characterize the somatic variant landscape of normal and cancerous cells, we generated a short-read WGS dataset spanning multiple cell types (Fig.1A, panel ii). Bulk samples included AML blasts, neutrophils, normal B-cells and T-cells, and cultured normal mesenchymal stromal cells (MSCs) (mean read depth 30x), all isolated with FACS; clonally expanded samples were derived from hematopoietic stem cells (HSCs) and multipotent progenitors (MPPs) with a mean read depth of 15x; single-cell WGS (scWGS) was performed on individual blasts and multiple mature and progenitor cell types (mean read depth 15x) via primary template-directed amplification (PTA) of the whole genome [20]. Importantly, phenotypically normal neutrophils (Fig.S1) were found to harbor the same AML driver variants as the blast population, indicating a shared clonal origin with the cell that eventually gave rise to the AML blast lineage and providing a phenotypically independent validation of the somatic SV calls.

As Nanopore data is known to exhibit non-random error patterns [21], we performed read correction and assembly with the state-of-the-art assembler hifiasm ONT (see Methods section *Nanopore data basecalling, read correction and genome assembly* and Fig.S2). We combined linear hg38 reference with two haplotype-resolved patient genome assemblies (HAP1 and HAP2) and constructed the pangenome graph with Minigraph-Cactus (Fig.1A, panel iii). The patient assemblies added 36.3 Mb of sequence absent from hg38 and approximately 6 million variant nodes (Fig.S3), which make up 42.35% of the total graph’s nodes, providing a substantially expanded genomic space for variant discovery.

The reference bias, evaluated with biastools in bulk samples (Fig.1B, Fig.S4), decreased in the pangenome-based alignments, as shown by the fraction of the alternative allele at the allelic bias plot. This indicates that providing individual-specific alternative alleles effectively mitigates the alignment penalties typically imposed by a single linear reference.

In all samples, we observed a marginal decrease in the proportions of mapped, primary mapped and properly paired reads, and in the percentage of high-confidence reads with mapping quality (MAPQ) = 60 in pangenome-based alignments (Fig.1C). These differences are expected consequences of surjection, the projection of graph alignments back onto hg38 coordinates that standard variant callers require: sequences absent from the hg38 cannot be represented in those coordinates and are therefore not visible to standard quality assessment tools [10].

A striking difference was shown for the number of reads with mate mapped to a different chromosome (MAPQ > 5), which decreased 3,316-fold (5425781±1933244 vs 1636.10±611.48) in pangenome-based alignments (Fig.1C, panel iii). To understand this effect, we traced high-quality split reads from hg38-based alignments of five bulk samples through the graph alignments to the surjected alignments (MAPQ > 55, to account for the drop of MAPQ caused by surjection; Fig.1D; and Methods: *Split reads tracing* section). After surjection, 28.55%±3.38% of these reads were rescued as properly mapped, mostly falling into centromeric, pericentromeric, and repeat-rich clusters of chromosomes 3, 17, and 21 (Fig.S5), consistent with the inclusion of pericentromeric and repeat-rich sequence in the patient assemblies that is absent or collapsed in hg38. However, reads spanning fully patient-specific graph nodes became unmapped after surjection because their sequence was absent from hg38. The remaining 70.35% ± 3.52% of the traced reads remained split, with one mate properly mapped and the other showing a low identity score in both graph-based and surjected alignments, making them promising candidates for SV discovery.

Categories include: ***hg38_interchromosomal*** - high-quality paired-end reads flagged by the hg38-based pipeline; ***Both_Reference*** - both reads in the pair align completely within reference genome paths inside the pangenome graph; ***One_Reference_One_LowID*** - one mate matches an hg38 path while its partner maps to a lower identity or structurally divergent path within the pangenome graph; ***One_Reference_One_Donor_Specific*** - one mate aligns to an hg38 path while its partner maps across the patient-specific node; ***Both_Donor_Specific*** - both reads are mapped across the patient-specific node; ***Both_Low_ID, Donor_Specific_Mixed_LowId*** - low-confidence mapping on the graph; ***Surject_both_mapped*** - both mates are successfully paired and mapped; ***Surject_mapped_one_only*** - only one mate is mapped, the second one fails; ***Surject_neither_mapped*** - both mates failed to be mapped.

### Individual-specific pangenome reference reduces the number of somatic indel calls, mainly in repeats

To assess how alignment on the pangenome affects the mapping quality of short variants, we compared somatic variant calls across bulk WGS samples (Fig.2A, Fig.S6). The pangenome-based analysis rescued 2.81±0.65% of variants that had lower mapping quality (MQ) in hg38-based alignments, producing a longer tail of the MQ distribution for the AML blast and neutrophil samples, but not for the B-cell sample (the T-cell sample was excluded from the analysis due to the overall low number of detected variants). This observation indicates that the pangenome improves the mapping quality of short variant calls.

**Figure 2:**
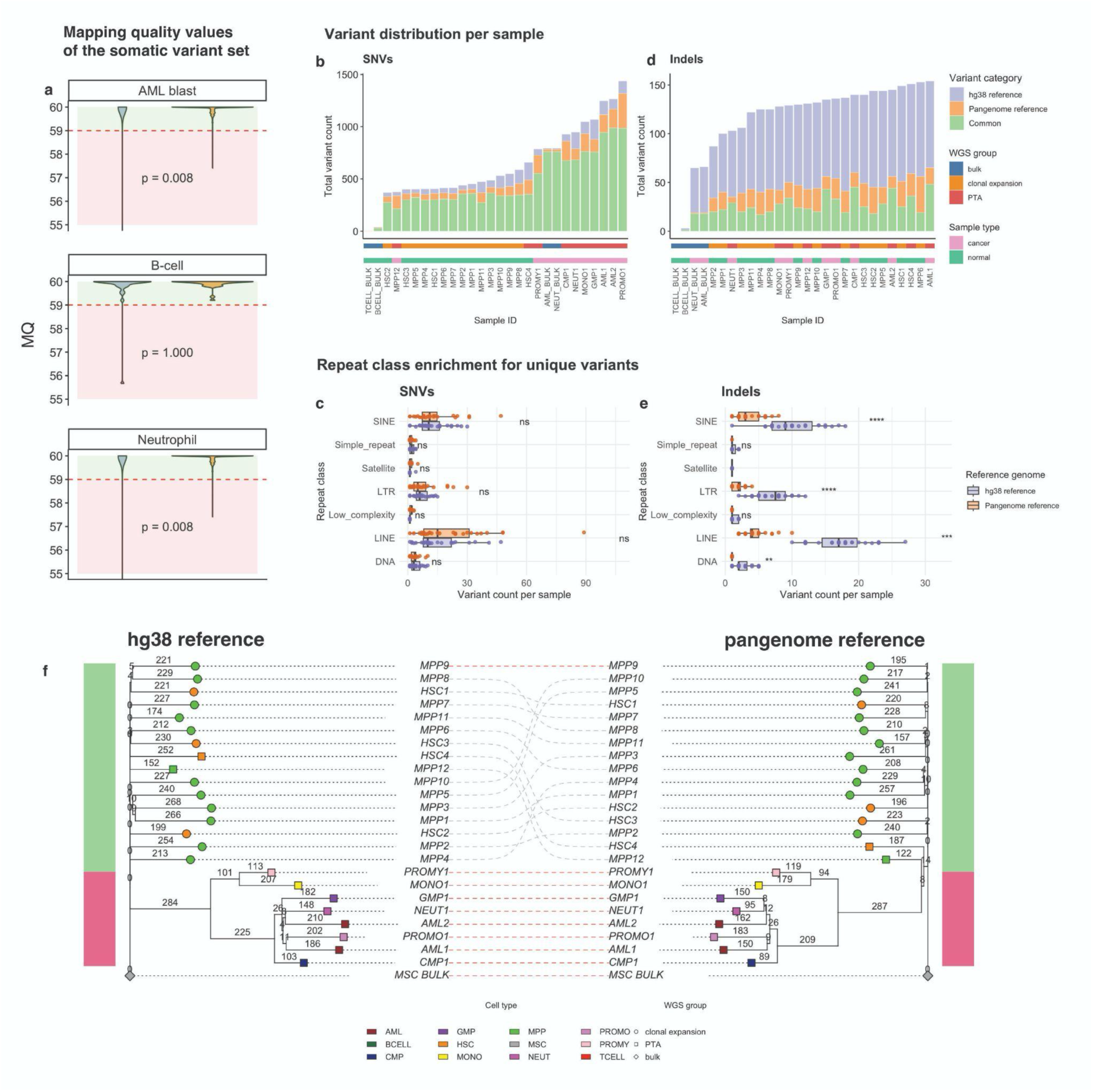
Somatic short variants called with hg38 and with the individual-specific pangenome. **(A)** Mapping quality (MQ) comparison for short variants called from hg38-based and pangenome-based alignments. The green zone indicates the variants retained with a MQ cutoff = 59; the red zone indicates the variants filtered out with the “BadMQ” tag. Violins are colored by the genomic reference used for mapping (violet - pangenome; orange - hg38). **(B)** Distribution of somatic SNVs per sample, ordered by total variant count. Bars are colored by variant categories: shared between hg38- based and pangenome-based alignments (common), unique to hg38-based alignments (hg38 reference), and unique to pangenome-based alignments (pangenome reference). **(C)** SNVs unique to hg38-based or to pangenome-based alignments located in repeat elements across all samples, stratified by repeat class. Boxplots represent variant counts in all samples in hg38-based and pangenome-based alignments. SINE - short interspersed nuclear element; LINE - long interspersed nuclear element; LTR - long terminal repeat; DNA - DNA transposon. **(D)** Distribution of somatic indels per sample, ordered by total variant count, stratified by the same categories as in B. **(E)** Indels unique to hg38-based or to pangenome-based alignments located in repeat elements across all samples, stratified by repeat class and by the same categories as in D. **(F)** SNV-based phylogenetic trees from hg38- based (left) and pangenome-based (right) alignments. The normal cells branch is green, the cancer cells branch is red.

Most SNVs were common with 83.5±9.85% for the hg38-based and 80.77±13.04% for the pangenome-based calls (Fig.2B). There were no significant differences in unique SNV numbers between the normal or cancerous cells, as well as between the bulk, clonally expanded or amplified with PTA samples (Fig.S7A, Fig.S7B). Pangenome-based alignments had a higher number of SNVs located in repetitive sequences overall, but the difference was not significant for any repeat class (Fig.2C, Fig.S7C).

In contrast to SNVs, indels unique to the pangenome were 58.73%±5.54% fewer than those unique to hg38 (Mann-Whitney U test, p-value = 4.224e-10), independently of biological condition (Fig.2D, Fig.S8A, Fig.S8B). This reduction likely reflects that the graph contains the patient-specific paths representing repetitive regions; reads from these regions align without the gaps that would produce false-positive indel calls. This is reflected in the repetitive sequences plot (Fig.2E, Fig.S8C) showing the overall and sample-specific reduction of indel variants located in short and long interspersed nuclear elements, long terminal repeats, and DNA transposons.

Single-cell WGS data obtained with PTA usually carries a substantial portion of SNVs introduced by strand displacement polymerases comprising mutational signature resembling SBS30 (Fig.S9A) [22]. We removed these artifacts with our previously described machine learning approach, PTATO [23], and compared the resulting mutational profiles of hg38-based and pangenome-based calls with the PTA artifact signature. In both cases, the profiles showed low cosine similarity to the artifact signature (Fig.S9B), allowing us to use both call sets for further analysis.

Finally, we compared somatic SNV-based phylogenetic trees of clonally expanded and PTA-derived samples aligned to hg38 and to the pangenome (Fig.2F). In both trees, normal HSCs and MPPs formed individual branches, consistent with polyclonality and differentiation trajectory in the hematopoietic system, whereas mature cells and AML blasts formed one distinct common branch [24]. PTA-derived and clonally expanded HSCs and MPPs were intermixed, indicating that the amplification method was not a confounder in the tree structure.

The pangenome-based tree was notably more balanced and symmetric, with a 40.4% drop in the Colless imbalance index (28 vs 47) and no sequential branching ladders (0 vs 4) in comparison to the hg38-based tree. The pangenome-based tree suggests more simultaneous, parallel diversification of subclones, rather than the prolonged sequential branches of the hg38-based tree. This is further supported by a higher number of small clades (10 vs 7 cherries and 5 vs 2 pitchforks, respectively) and a higher treeness parameter (0.135 vs 0.123), indicating a more shared evolutionary path between individual samples. No ground-truth phylogeny exists for this patient, so we cannot establish which topology is correct.

### Somatic SV calling with the individual-specific pangenome

To determine how mapping against the individual-specific pangenome affects SV calling, we compared somatic SVs identified in hg38-based and pangenome-based alignments of bulk WGS samples. Pangenome-based alignments resulted in more SVs in both normal and cancerous samples compared to hg38-based alignments, with the most pronounced differences in deletions (7 vs 1) and duplications (4 vs 2) (Fig.3A, Fig.3B, Fig.S10, Fig.S11). Both hg38-based and pangenome-based analyses detected the two diagnostic driver events, *NUP98-NSD1* and *FLT3*-ITD, in the bulk AML and neutrophil samples, so the additional pangenome SVs did not displace the established driver calls (Fig.3B, panels ii and iii).

**Figure 3:**
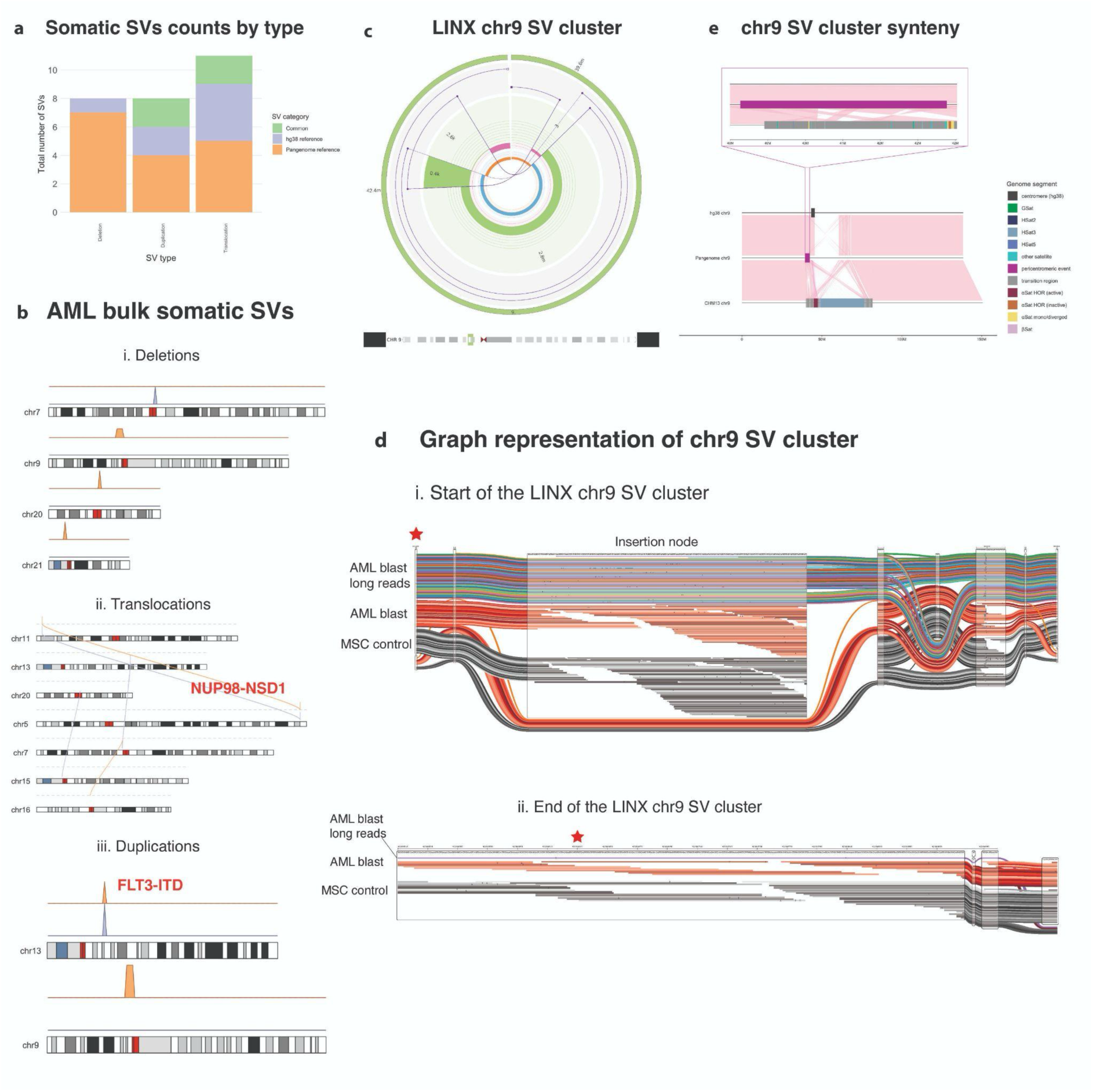
Somatic SVs called with hg38 and with the individual-specific pangenome. **(A)** Somatic SV counts per bulk WGS sample, stratified: shared between hg38-based and pangenome-based alignments and unique per alignment category. **(B)** Ideograms of somatic SV events in the AML bulk sample identified with pangenome-based and hg38-based analyses across deletions (i), translocations (ii), and duplications (iii). Driver SV events are shown in red. **(C)** Circos plot of the pericentromeric SV cluster at 9p12, initially called as somatic in pangenome-based alignments and subsequently resolved as germline. **(D)** SequenceTubeMap graph alignment visualization of the pericentromeric SV starting breakend in AML sample long-read data (top, colorful), AML bulk short-read data (middle, red), and MSC control short-read data (bottom, gray) samples for the start (i) and end (ii) breakends of the SV. Red stars mark the exact positions of the breakends. **(E)** Synteny plot of pairwise alignments of chromosome 9 (hg38 on pangenome and pangenome on T2T-CHM13).

The most distinct pangenome-specific somatic SV we identified spanned 2.8 Mb in the pericentromeric region 9p12. It was represented by multiple insertion/deletion events along with rearrangements of highly homologous sequences and a 400 bp duplication toward the centromere (Fig.3C). To further investigate whether this SV was somatic or germline, we examined high-quality graph alignments of short and long reads of the AML bulk sample and the corresponding MSC control sample. The split reads supporting the breakend (Fig.3D, panel i) mapped to a 294 bp graph node in reverse orientation, within a low-complexity polyT region. Moreover, the reads mapped to this node were also present in the short-read data of the AML bulk sample and of the MSC control, and in the AML bulk long-read data, indicating that this variation is a germline insertion. The second breakend showed a similar insertion pattern for both cancer and normal samples, although with low support from the AML bulk long-read data (Fig.3D, panel ii).

To understand the reason behind the misclassification of this SV, we looked at its GRIDSS output tags. The breakpoint of this event was anchored within a 300 bp repetitive sequence tract (inexact homology tag), and GRIDSS reported the same junction as multiple overlapping calls. Although manual inspection showed the presence of reads supporting this junction in the MSC control sample, the repetitive sequence at the breakpoint likely prevented their assembly into a split-read call, reducing the reported normal support to zero after somatic filtering.

Analysis of the insertion-related nodes’ sequences with the web version of MegaBLAST (date of accession: 14.07.2026, [25]) showed that the sequences of both insertion nodes match BAC clone CH17-329N15 from chromosome 9. The node at the first breakend contains a microsatellite motif (AC)n, and the node at the second breakend contains a 161 bp sequence that maps twice in tandem on the BAC clone and is consistent with alpha-satellite sequence. These observations indicate that the nodes derive from a pericentromeric locus and that the pangenome placed these reads correctly. Synteny analysis of the chromosome 9 sequences between hg38, the individual-specific pangenome, and the T2T-CHM13 reference, annotated with the map of human pericentromeres and centromeres [26], also placed the SV within a pericentromeric transition region adjacent to chr9 transition and satellite sequences, supporting the origin of these nodes (Fig.3E).

Manual inspection of SV breakpoints in IGV (Additional file 2) and PCR amplification of the junction fragments in AML bulk, neutrophil bulk and MSC bulk samples (Fig.S12) showed that only *FLT3-*ITD and *NUP98-NSD1* were specific for the samples of tumor origin. All other somatic SVs reported by GRIDSS were germline variants or technical artifacts of the SV calling process in centromeric regions. This case illustrates that current somatic SV callers, which work in hg38 coordinates, cannot fully resolve the graph-to-linear representation of structural variation. SVs called with such tools from pangenome-based alignments require inspection at the raw-read level, especially in pericentromeric and repeat-rich regions, before biological or clinical interpretation.

### The individual-specific pangenome removes artifactual candidate driver calls but can introduce new ones

To test whether the use of the pangenome changes the driver landscape, we compared candidate driver calls from hg38-based and pangenome-based alignments in bulk and scWGS data (Fig.4A). The pangenome gave fewer candidate driver calls, mainly due to a significant drop in frameshift insertions upstream of polyG and polyC repeats in PTA-derived scWGS samples (7.92±4.94 vs 1.15±1.14, p=0.0012, Fig.S13). Almost all single cells, except for one MPP cell, shared the *IDH2* missense driver with the bulk samples of the cancerous origin, but lacked the *WT1* and *FLT3* drivers, consistent with the overall low coverage of the regions where these drivers are located despite the overall presence of reads supporting them (Fig.S14).

**Figure 4:**
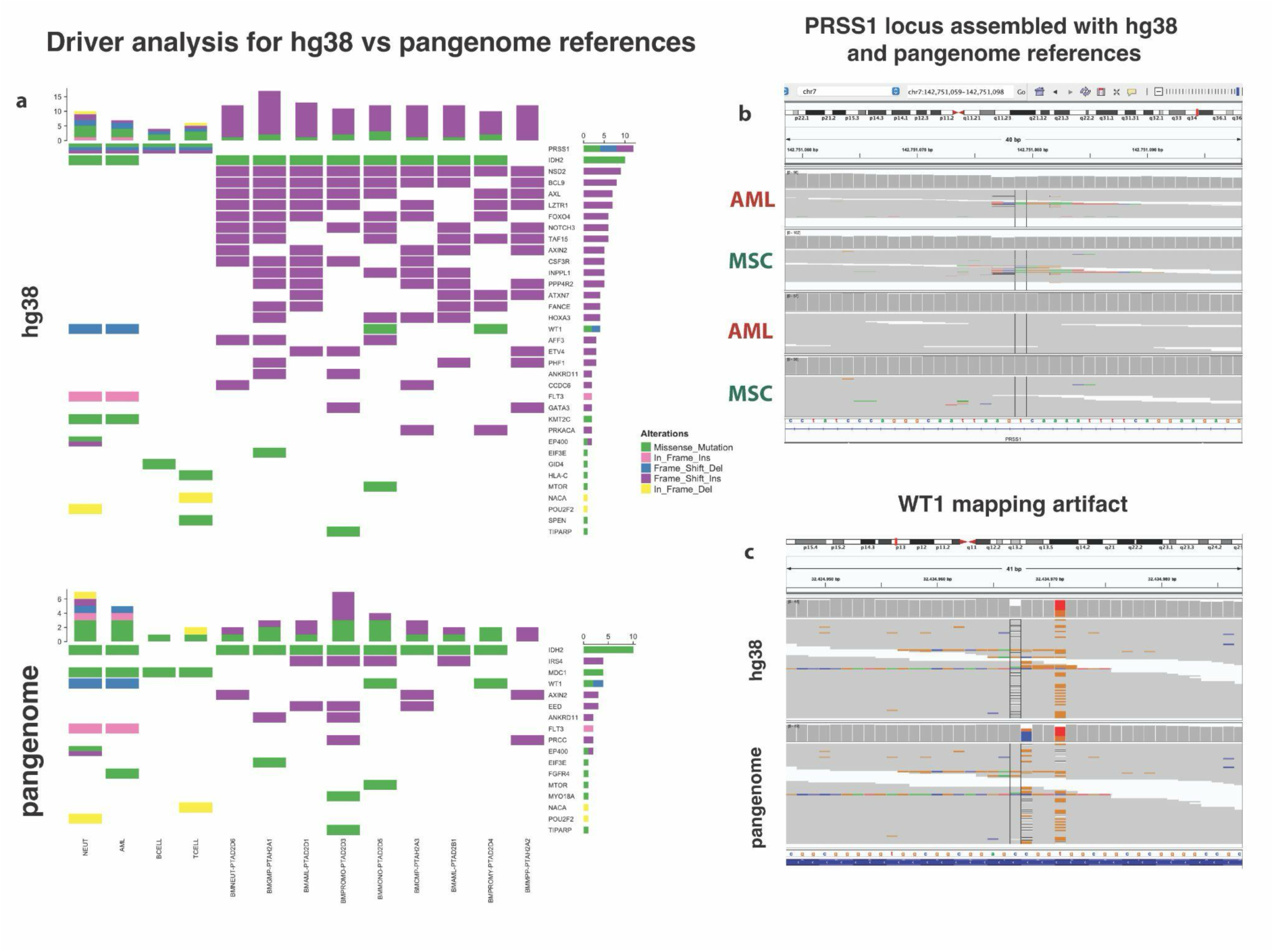
Candidate driver variants called with hg38 and with the individual-specific pangenome. **(A)** Driver analysis results comparison of hg38-based set of variants (upper panel) and pangenome-based set of variants (lower panel). Cell colors correspond to the mutational type. **(B)** IGV alignment view of the *PRSS1* locus comparing hg38-based (top two tracks) and pangenome-based (bottom two tracks) alignments in bulk AML and MSC control samples. **(C)** IGV alignment view of indel artifact of graph-based mapping in *WT1* gene, bulk AML sample.

Three variants with high or moderate predicted impact on an alternative isoform of the *PRSS1* gene (ENST00000486171) were called with hg38 but not with the pangenome. We validated this result by manual inspection in IGV, which revealed substantially reduced alignment noise across the extended *PRSS1* region in pangenome-based alignments (Fig. 4B). Sanger sequencing of this region showed no deviation from hg38 (Fig.S15A), confirming that the three *PRSS1* calls were hg38 alignment artifacts that the pangenome resolved.

Despite the observed reduction in false positive calls, pangenome-based alignments can still generate indel-based artifacts. We identified a novel artifact of graph-based alignment and surjection in a low-complexity region in exon 1 of the *WT1* gene (chr11:32,434,946-32,434,986, Fig. 4C). In hg38-based alignments, this region carried a 1 bp indel at the CC site (heterozygous C deletion) and a T/G missense SNV at position 3 of the GGTGG motif. In contrast, in the surjected pangenome-based alignments, the indel calls were split between position +1 and the T/G SNV, practically erasing the SNV from the filtered output and from driver analysis results. Validation via Sanger sequencing (Fig.S15B) revised the indel to a homozygous C insertion in the polyG tract (chr11:32,434,951-32,434,952) and relocated the SNV to chr11:32,434,964 (C>G change), so neither hg38-based nor pangenome-based calls matched the validated genotype. This example shows that indels in low-complexity regions need additional verification, particularly when the pangenome graph introduces alternative path representations at sites of known somatic relevance.

## Discussion

Since its release in 2013, hg38 has been established as the widely accepted genome reference. It was used as the universal “ground truth” to evaluate the mutational landscape of countless samples and to define what is considered detectable in most bioinformatics tools and clinical frameworks [27,28]. The T2T-CHM13 assembly [29] added the missing 8% of the genome, nearly 200 Mb of sequence, but remained a linear, “one-size-fits-all” reference.

The individual-specific pangenomic approach is the logical continuation of our understanding of the genome organization at the patient level. Paired tumor-normal callers have already been run on population pangenome-based alignments projected to a linear reference and on personalized diploid assemblies, but not on short-read bulk and single-cell alignments surjected from an individual-specific pangenome [3]. Our work highlights the concrete steps of the analysis that require additional checks because of the current gaps in compatibility between graph alignments and downstream tools built for linear references.

Because standard somatic callers and annotation tools need linear coordinates to work with, downstream analyses heavily rely on surjection to hg38 reference. This allowed us to rescue 28.55% of the reads split in hg38-based alignments, but the reads in fully patient-specific nodes were dropped, making somatic variants located in these nodes impossible to analyze. Somatic callers that work directly with the graph genome would be able to access these variants, but they are still limited: Pansoma, a deep learning caller similar in design to DeepVariant and DeepSomatic [30,31], works only in tumor-only mode [32], and paired tumor-normal calling on the graph exists only for SVs, by subtracting normal calls from tumor ones [33], and for small variants it remains an open challenge for the field.

The consequences of such inability to fully accommodate the somatic variation along with the personal one became the source of misleading SV calling. The false-positive somatic SV in the pericentromeric region of chromosome 9 arose because GRIDSS, working in hg38 coordinates, reported a germline junction within a repeat as multiple overlapping calls and did not assemble the supporting reads in the normal sample. This example shows that appropriate assembly and mapping of a highly repetitive region do not automatically translate into improvements in the subsequent variant calling and interpretation. Manual inspection of graph alignments at the breakends, together with PCR, resolved this case, but this approach cannot be scaled to all SV calls in any given sample. Nevertheless, the pangenome placed reads in this pericentromeric region correctly, showing patient-level variation in regions that hg38- based analyses cannot properly assess.

For short variants, indels unique to the pangenome were 58.73% fewer than those unique to hg38. Most of them were located in LINE, SINE and LTR elements, which are hard to map with short reads because reads shorter than a repeat cannot be placed uniquely among nearly identical copies [34,35]. Interestingly, we also saw the same effect for the *PRSS1* gene, which is notoriously difficult to map with short reads because of its homology with the *PRSS2* gene and the *PRSS3P2* and *TRY7* pseudogenes. The current hg38 build includes a common deletion polymorphism of both pseudogenes, allowing the reads coming from this part of the genome to be attributed to the *PRSS1* [36,37]. By resolving the location of the reads spanning these loci, pangenome-based alignments removed the cluster of false-positive variants. This finding is particularly relevant because common germline *PRSS1-PRSS2* variants are linked to asparaginase-associated pancreatitis in children treated for acute lymphoblastic leukemia, and NGS mismapping at *PRSS1* has produced false-positive calls in chronic pancreatitis testing [38–40]. Conversely, surjection of pangenome-based alignments split a driver indel call in the low-complexity exon 1 of *WT1*, and hg38-based short-read calls did not match Sanger sequencing at this site either, likely due to being error-prone in homopolymers and low-complexity sequences [41,42]. Correct *WT1* calling is clinically relevant, because additional *WT1* mutations were observed in a subgroup of patients with *NUP98-NSD1* and *FLT3*-ITD in small pAML cohorts with the very poor outcome [18,43]. Graph-based results, like any other, therefore require verification at clinically relevant loci.

The challenges of variant calling are further amplified in scWGS data. PTA reduces allelic dropout and skewing compared with other whole-genome amplification methods [20], but the data still contain regions with uneven coverage and amplification artifacts, which the PTATO model separates from true base substitutions with the accuracy of 0.74 [23]. Alignment to hg38 can add mapping artifacts to this noise, increasing the risk that true somatic variants are indistinguishable from it or lost entirely, limiting the use of scWGS for reconstructing clonal architecture in healthy tissue and cancer. In our data, the pangenome removed frameshift insertion artifacts next to polyG and polyC repeats in single cells, which reduced false-positive candidate driver calls. The use of the pangenome reference also changed the inferred phylogeny of SNV-based tree, although without a ground truth we cannot tell which tree is more accurate.

Our study has several limitations. We analyzed a single patient, so the size of the observed effects may differ in other patients and cancer types. Orthogonal validation covered selected loci (somatic SV calls inspected via PCR, *PRSS1* and *WT1* loci inspected by Sanger sequencing), but not the genome-wide differences in SNV and indel calls. Because no graph-native caller supports paired tumor-normal small variants calling at the moment, we used standard linear-reference callers and downstream analysis tools on surjected alignments, which cannot assess variants in patient-specific sequence. Finally, the pangenome was built from 30x Nanopore Simplex reads, and more accurate or complete assemblies may represent repetitive regions better.

## Conclusions

In the described pediatric AML case an individual-specific pangenome removed false-positive indels mainly in repetitive and polyG/polyC regions, and eliminated the false-positive drivers in a highly homologous *PRSS1* locus. However, several problems remain: surjection to hg38 split a *WT1* driver call, novel somatic SVs called with the pangenome turned out to be germline variants or artifacts, and graph-native paired callers for somatic small variants are still missing. Overall, individual-specific pangenomes can reduce artifact calls, especially in scWGS data, but they have yet to reach the maturity required for routine clinical use, and pangenome-based calls need read-level inspection and orthogonal validation.

## Methods

### Patient samples

Bone marrow samples of the patient at the AML diagnosis timepoint were obtained via the Biobank of the Princess Máxima Center for Pediatric Oncology, with ethical approval under proposals PMCLAB2018.007 and PMCLAB2021.265 in accordance with the Declaration of Helsinki. Part of the data for the researched patient (11 clonally expanded MPPs, 3 clonally expanded HSCs, as well as bulk AML blast and MSC samples) was previously reported by Brandsma et al [24].

### FACS and HSPC culture

Bone marrow mononuclear cells were stained for FACS after thawing. We defined AML blasts based on diagnostic immunophenotyping data, and these were defined as CD33, CD38, and CD117 positive, and CD34 dim. At least one of these markers was used for gating of normal cells, so that these were negative for the blast marker. Hematopoietic stem cells (HSCs) were identified using the following surface markers: Lin−CD11c−CD16−CD34+CD38−CD45RA−CD90+. Multipotent progenitor cells (MPPs) were Lin−CD11c−CD16−CD34+CD38−CD45RA-CD90-; common myeloid progenitors (CMPs) were Lin−CD11c−CD16−CD34+CD38+CD45RA− and granulocyte-macrophage progenitors (GMPs) were Lin−CD11c−CD16−CD34+CD38+/CD45RA+.

Mature cell types were sorted using a separate antibody panel, also including a blast marker for negative gating. B-cells were identified as CD34-CD45+CD20+, monocytes were CD34-CD45+CD14+, and megakaryocyte progenitors were CD34-CD45- CD42b+. Promyelocytes were identified as CD117+HLADR-, neutrophils as CD117- HLADR-CD11b+CD14-, promonocytes as CD117-HLADR+CD15dimCD14-, and monocytes as CD117-HLADR+CD15+CD14+.

Different cell populations were purified on an SH800S Cell Sorter (Sony). First, blasts were sorted in bulk for DNA isolation. Single cells of blasts and all other populations were sorted in a 96-well plate prepared with PTA cell buffer and stored at -80°C. Additionally, HSPCs were index sorted in a flat-bottom, 384-well plate prepared with 75 μL HSPC culture medium per well, or sorted in bulk directly in a 6-well plate (Sarstedt) with 2 mL pre-warmed HSPC culture medium per well. HSPC culture medium consisted of StemSpan SFEM II medium (Cat. No. 09655; Stem Cell Technologies) supplemented with SCF (100 ng/mL, Cat. No. 130-096-692; Miltenyi), FLT3 ligand (100 ng/mL, Cat. No. 130-096-474; Miltenyi), IL-6 (20 ng/mL, Cat. No. 130- 093-929; Miltenyi), IL-3 (10 ng/mL, Cat. No. 130-093-909; Miltenyi), TPO (50 ng/mL, Cat. No. 130-095-745; Miltenyi), UM729 (500 nM, Cat. No. 72332; Stem Cell Technologies) and StemRegenin-1 (750 nM, Cat. No. 72342; Stem Cell Technologies).

HSPCs sorted in HSPC culture medium were clonally expanded (for single-cell WGS) and cultured in bulk (for subsequent Nanopore sequencing) for 4-7 weeks at 37°C, 5% CO2, 20% O2 before harvesting. Mesenchymal stromal cells (MSCs) were cultured from a bone marrow fraction of 500,000 cells/well in 12-well culture dishes with DMEM- F12 medium (GIBCO), supplemented with 10% FCS. Medium was refreshed every other day to remove non-adherent cells, and MSCs were harvested when confluent, after approximately 2-3 weeks.

### FACS antibodies

All antibodies were obtained from BioLegend, except for CD117-BV421 (BD Bioscience). For AML blasts and HSPCs the following antibodies were used: CD34- BV421 (clone 561, 343609), CD38-PE (clone HIT2, 303505), CD45RA-PerCP/Cy5.5 (clone HI100, 304121), CD90-APC (clone 5E10, 328113), Lineage (CD3/CD14/CD19/CD20/CD56)-FITC (clones UCHT1, HCD14, HIB19, 2H7, HCD56, 348701), CD11c-FITC (clone 3.9, 301603), CD16-FITC (clone 3G8, 302005), CD33- PE/Cy7 (clone WM53, 303433), CD13-APC (clone WM15, 301705) and CD117- PE/Cy7 (clone 104D2, 313211).

For mature populations the following antibodies were used: HLADR-FITC (clone L243, 307603), CD14-AF700 (HCD14, 325614), CD20-FITC (clone 2H7, 302303), CD42b- PE (clone HIP1, 303905), CD34-BV421 (clone 561, 343609), CD117-PE/Cy7 (clone 104D2, 313211), CD117-BV421 (clone YB5.B8, 562434, BD Bioscience), CD33- PE/Cy7 (clone WM53, 303433), CD45-APC (clone 2D1, 368511), CD11b-APC (clone ICRF44, 301309), CD15-PE (clone HI98, 301905).

### DNA isolation and whole-genome sequencing

DNA was isolated from cell pellets of blasts, MSCs, B-cells, T-cells, neutrophils and clonally expanded HSCs and MPPs using the DNeasy DNA Micro Kit (Qiagen), following the manufacturer’s instructions. The standard protocol was slightly adjusted by adding 2 µL RNase A (Qiagen) during the lysis step and eluting DNA in 50 µL low EDTA TE buffer (10 mM Tris, 0.1 mM EDTA, G Biosciences). DNA libraries for Nanopore sequencing (cultured HSPCs and sorted AML blasts) were prepared with Ligation Sequencing Kit V14 (Oxford Nanopore Technologies) according to the manufacturer’s instructions. The libraries were sequenced using PromethION with R10.4.1 flow cells at a depth of 30x for HSPCs and 15x for AML blasts. DNA from single cells was isolated and amplified using the ResolveDNA™ Whole Genome Amplification Kit (BioSkryb) version 1 according to the manufacturer’s instructions. DNA libraries for Illumina sequencing were generated from at least 45 ng genomic DNA using standard protocols. For PTA-amplified DNA, at least 500 ng genomic DNA was used. The libraries were sequenced using Illumina Novaseq 6000 (2x150bp) at a depth of 15x (clonally expanded and single cells) and 30x (bulk samples).

### PCR and Sanger sequencing for SVs and short variants validation

Primers for SV and short variants were designed with the Primer3 1.1.3 tool [44]. For SVs the output of GRIDSS with the 50 bp offset was used. The DNA amplification from bulk samples for selected loci validation only was performed via PTA. The PCR reaction was performed for 35 (for SVs) or 40 (for short variants) cycles on a T100 Thermal Cycler (Biorad). 5 µL of PCR products were loaded on a 1% agarose gel to verify the amplification of the targeted region. 2 µL of PCR products were mixed with 8 µL of 10 µM sequencing primer and used for Sanger sequencing at Macrogen. The correct sequence amplification and the presence of SNVs and indels were verified using Benchling (date of accession: 08.09.2026). Sanger sequencing was performed on 12 ng of isolated genomic DNA using the GoTaq G2 colorless mastermix (Promega; M7832) following the manufacturer’s instructions. Detailed PCR parameters and primers to isolate SV/variant-containing loci can be found in Additional file 3.

### Pangenome-based workflow

#### Nanopore data basecalling, read correction and genome assembly

Basecalling of raw Nanopore WGS data was performed with dorado v0.8.0 [45] using the super accurate model (version dna_r10.4.1_e8.2_400bps_sup@v5.0.0). Read correction was performed with hifiasm --ont [21] and with the dorado correct function with HERRO algorithm [46], and the resulting assemblies were compared (Fig.S2). To assemble the genome *de novo*, in both of the cases we used hifiasm version 0.24.0- r703 [47] with default parameters for assembling ONT reads. The hifiasm ONT assemblies were used for pangenome construction.

We evaluated the assembly quality (both haplotype-resolved and primary) using gfastats v1.3.6 [48] by comparing L50, N50, total number of contigs, lengths of the shortest and longest scaffolds, and the total length of the assembled genome. Additionally, we performed k-mer-based evaluation of the genome assemblies with Merqury v.1.3 [49] using short read data of two bulk samples (T-cell and MSC). We estimated the assemblies’ completeness using compleasm v.0.2.7 [50] with the BUSCO primates lineage. Finally, we performed pairwise alignment of our assemblies against T2T-CHM13v2.0 [51] with minimap2 [52] and visualized the chromosome-level coverage using the modified ‘plot_coverage’ function from the pafr package in R [53]. The reference bias of genomic alignments was assessed with biastools v0.3.0 [54].

#### Personal pangenome graph construction

Individual-specific pangenome graph was constructed from haploid Hifiasm assemblies HAP1 and HAP2 and canonical GRCh38.p14 (hg38) genome with Minigraph-Cactus v2.9.3 [55] using corresponding parameters: “--reference hg38 -- filter 1 --haplo --giraffe filter --viz --odgi --chrom-vg clip filter --chrom-og --gbz clip filter full --gfa clip full --vcf --vcfReference hg38” to produce the necessary indexes, chromosome-level visualizations and odgi-compatible representation of the pangenome.

#### Alignment and visualization of short-read data on pangenome graph

To produce sample-level alignments on an individual-specific pangenome graph, we used the haplotype-aware vg v1.72.0 Giraffe aligner [56] with a GBZ-formatted graph and XG index as inputs and GAM and surjected BAM files as outputs. Surjected BAM files were processed with the same standard workflow as hg38-based alignments. Visualization of the regions of interest was performed by extracting a target subgraph using vg find. The graph was sliced along the hg38 path coordinates of interest. We included two degrees of extra node context (-c 2) to preserve the surrounding graph topology at the boundaries. The resulting subgraph was subsequently visualized using sequenceTubeMap [57].

#### BAM files comparison

BAM file statistics were obtained with samtools v.1.17 flagstat and samtools stats commands [58].

#### Split reads tracing

High-confidence split and discordant reads (MAPQ > 55, mate is mapped to another chromosome) were isolated from hg38-based alignments and traced through GAM files and surjected linear BAM files. Read fragments were stratified by genomic origin and donor specificity in the GAM file, separating reference-spanning sequences from donor-specific, mixed, and low-identity reads. Following surjection, the mapping of both mates within each fragment pair was evaluated. Surjected BAM fragments were classified into three categories based on alignment status: Surject_both_mapped, where both read mates successfully resolved, properly paired, and mapped within linear coordinates; Surject_mapped_one_only, indicating an asymmetrical alignment where a single mate dropped out or remained split/unmapped; and Surject_neither_mapped, where both reads failed to be mapped. The final percentages of each category were quantified, normalized, and visualized using a Sankey diagram with the networkD3 package in R to illustrate the efficiency of the pangenome graph mapping in rescuing split reads.

### Standard workflow

#### Read mapping

WGS data analysis was performed using an in-house developed ASAP pipeline [59].

Sequencing reads were first mapped to hg38 using bwa-mem2 v2.2.1 [60] with settings "-M -c 100", then duplicates were marked using GATK [61] MarkDuplicatesSpark and base recalibration was performed using GATK’s BaseRecalibration. All GATK tools here were from version 4.4.0.0.

#### Mutation calling, filtering, and annotation

Mutation calling was performed using best practices in GATK’s variant calling. We used HaplotypeCaller per sample with options “-ERC GVCF” and consolidated files in GVCF format using GenomicsDBImport with options “--genomicsdb-shared-posixfs- optimizations true --bypass-feature-reader”. Next, GATK’s genotyping was performed with GenotypeGVCFs, and variants filtering was implemented using VariantRecalibrator (for indels we used “-an QD -an MQ -an MQRankSum -an ReadPosRankSum -an FS -an SOR -mode INDEL” settings, and for SNPs “-an QD - an MQ -an MQRankSum -an ReadPosRankSum -an FS -an SOR -mode SNP”) and ApplyVQSR (with parameters for indels “--truth-sensitivity-filter-level 99.9 -mode INDEL” and for SNPs “--truth-sensitivity-filter-level 99.9 -mode SNP”).

GATK VariantFiltration settings were given as:

> “--filter-expression ’QD < 2.0’ --filter-expression ’MQ < 40.0’ --filter-expression ’FS > 60.0’ --filter-expression ’HaplotypeScore > 13.0’ --filter-expression ’MQRankSum < - 12.5’ --filter-expression ’ReadPosRankSum < -8.0’ --filter-expression ’MQ0 >= 4 && ((MQ0 / (1.0 * DP)) > 0.1)’ --filter-expression ’DP < 5’ --filter-expression ’QUAL < 30’ -- filter-expression ’QUAL >= 30.0 && QUAL < 50.0’ --filter-expression ’SOR > 4.0’ --filter-name ’SNP_LowQualityDepth’ --filter-name ’SNP_MappingQuality’ --filter-name ’SNP_StrandBias’ --filter-name ’SNP_HaplotypeScoreHigh’ --filter-name ’SNP_MQRankSumLow’ --filter-name ’SNP_ReadPosRankSumLow’ --filter-name ’SNP_HardToValidate’ --filter-name ’SNP_LowCoverage’ --filter-name ’SNP_VeryLowQual’ --filter-name ’SNP_LowQual’ --filter-name ’SNP_SOR’ -cluster 3 -window 10”

Callable regions filtering was done using GATK 3.8 CallableLoci with options ‘-- minBaseQuality 10 --minMappingQuality 10 --minDepth 8 --minDepthForLowMAPQ 10’ for PTA samples and ‘--minBaseQuality 10 --minMappingQuality 10 --minDepth 15 --minDepthForLowMAPQ 20’ for WGS samples. VCF files were subsetted by common callable regions across all samples in the batch.

Finally, annotation of variants was done with VEP v112.0 and settings “--vcf --plugin AlphaMissense,file --plugin NMD --plugin dbNSFP”.

#### Somatic mutation filtering for WGS samples

Somatic Mutations Rechecker and Filtering (SMuRF) v3.0.2 was used for obtaining high-confidence clonal somatic mutation calls [62]. These were mutations that (A) were positioned on autosomal chromosomes, (B) had a GATK phred-scaled quality score >= 100, (C) had a mapping quality of 59 (30x coverage) or 55 or higher (15x), (D) had a base coverage of at least 10 (30x) or 5 (15x), (E) had a GATK genotype quality of 99 (heterozygous) or 10 (homozygous) in the sample. Additional filtering included removal of variants found in a panel of normals, PON, which can be found in [63].

#### Mutation filtering for PTA single-cell WGS data

For PTA scWGS samples, we used the PTA Analysis TOolbox (PTATO) v1.3.3 [23] to separate PTA-induced artifacts from true mutations. Cosine similarity was computed with the cos_sim_matrix function from the MutationalPatterns package v.3.18.0 [64] in R.

#### Merging WGS and scWGS samples with HaplotypeCaller

To compile the clonally expanded WGS samples and scWGS samples, both sequenced with 15x coverage, we used the GATK HaplotypeCaller tool to re-evaluate variant calls via local re-assembly of haplotypes, effectively processing more than 15 samples at once, which our ASAP pipeline does not currently support. The output VCF file was filtered to contain only somatic SNVs and used as an input for phylogenetic tree building. Bulk samples (except for the germline control) were excluded from the tree building.

#### Structural variant analysis

SV analysis was performed using the SV calling module of ASAP, which integrates GRIDSS v.2.13.2, PURPLE v.3.7.1, LINX v.1.23.6, and AMBER v.3.9 tools [65–67].

GRIDSS VCF outputs filtered by PASS tag were annotated with AnnotSV v.3.5.8 [68]. All SVs with a maximum population allele frequency for benign gains or losses exceeding 1% were excluded from the downstream analysis. Graph alignments of the specific region of interest were extracted from the graph using the vg toolbox and visualized using sequenceTubeMap [57] for the reads with MAPQ > 55. Synteny plots were created based on the minimap2 v.2.28 pairwise alignment of chromosome 9 with the “-x asm10” parameter and plotted with the gggenomes v.1.1.3 package in R. Centromere annotations were plotted according to the repeat annotation workflow results from [26].

### Driver analysis

Potential driver variants were identified based on the framework from [69] and cancer gene list from OncoKB database [70]. In short, SNVs and indels were filtered against matched germline control, and only variants labeled with “PASS” tag after SMuRF, with “HIGH” or “MODERATE” tag by Variant Effect Predictor (VEP) [71] or “likely pathogenic" tag by AlphaMissense [72] were retained. Oncoplots were generated with the oncoPrint function from the ComplexHeatmap package v.2.24.1 in R.

### Phylogenetic tree construction

Phylogenetic reconstruction was performed on the GATK HaplotypeCaller output VCFs using CellPhy v0.9.2, which builds on RAxML-NG, a maximum-likelihood framework for phylogenetic inference [73,74]. CellPhy considers the allelic dropout rate and amplification errors per sample and constructs the most likely tree based on these estimates. CellPhy was run on the phred-scaled genotype likelihoods (PL) with standard settings, including the “GT16 + FO + E” model. CellPhy was also used to map mutations to the tree (using the “--mutmap” function) with the “--opt-branches off” setting. The CellPhy “--support” function was used to run 100 bootstrap replicates. The trees were rooted with treeio’s “root” function, which is a wrapper for the “root” function from the R package ‘ape’. Phylogenetic trees were plotted with the cophylo function from phytools v2.5-2 in R. Both phylogenies were assessed and compared with R package treestats v.1.70.8.

### Somatic variants comparison between standard and pangenome-based workflows

To compare variants called and filtered using the pangenome versus hg38, we overlaid short variants annotated by VQSR (Variant Quality Score Recalibration) from both approaches and compared their MQ and VQSR status. This allowed us to evaluate the pangenome’s ability to rescue poorly aligned hg38 variants and to assess potential pangenome-specific bias. Repeats were annotated with RepeatMasker hg38 track in UCSC Genome Browser [75].

### Statistical analysis and plotting

Group comparisons were performed with the Mann-Whitney test (unless otherwise specified) in R v.4.5.0 using the “wilcox.test” function, with Bonferroni correction for multiple comparisons where applicable. All mean values are given in the mean ± SD format, where SD is the standard deviation. Graph plotting is performed with ggplot2 v.4.0.2 in R.

## Additional files

Additional file 1.docx - This file contains Supplementary Figures S1-S15.

Additional file 2.pdf - This file contains IGV visualizations of SV breakends in hg38- based and pangenome-based alignments.

Additional file 3.xlsx - This file contains primers’ sequences and PCR protocols used for amplification of validated loci.

## Declarations

### Ethics approval and consent to participate

Bone marrow samples were obtained through the biobank of the Princess Máxima Center for pediatric oncology. The Medical Research Ethics Committee NedMec (METC NedMec, Utrecht, The Netherlands) determined that the biobank protocol does not fall within the scope of the Dutch Medical Research Involving Human Subjects Act (WMO) and is therefore not subject to the requirements of the WMO. The present study was subsequently reviewed and approved by the Biobank and Data Access Committee of the Princess Máxima Center (project approval numbers: PMCLAB2018.007 and PMCLAB2021.265). The patient and/or their parents or legal guardians provided written informed consent for the storage and future research use of the patient’s biological materials and associated data.

## Consent for publication

The patient and/or their parents or legal guardians provided written informed consent for the storage and future research use of the patient’s biological materials and associated data.

## Availability of data and materials

Short-read data (BAM format) has been deposited at the European Genome-phenome Archive (EGA) [76], which is hosted by the EBI and the CRG, under accession number EGAS50000002184.

Variant calls and synteny data are deposited in the Mendeley repository: Kulaeva, Elizaveta (2026), “Somatic variant detection using an individual-specific pangenome reference: a case study of a pediatric acute myeloid leukemia patient”, Mendeley Data, V1, doi: 10.17632/5f2bb4nfpw.1 [77].

The code associated with the publication is available on Github (https://github.com/ProjectsVanBox/individual-specific-pangenome) [78] and in Zenodo [79].

## Authors’ contributions

EK analyzed and interpreted the sequencing data, designed the split reads tracing workflow, implemented the code, wrote the initial draft (lead) and addressed the co-authors’ comments. EJMB performed FACS analysis, wrote part of the Methods section and addressed the co-authors’ comments. MvR set up the PTATO tool for the pangenome reference-based usage and contributed to the interpretation of the SV calling results. RH contributed to the analysis and interpretation of the sequencing data. LLMD performed cell culturing, FACS analysis, DNA extraction for the ONT WGS and substantially revised the manuscript. LT performed the short-read and ONT WGS library preparation. AMB performed FACS analysis. MV performed short-read and single-cell WGS library preparation. MB performed the PCR-based validation of the variants in *WT1* and *PRSS1* and substantially revised the manuscript. SM performed the single-cell WGS library preparation. PP contributed to the design of the study and revised the manuscript. RvB contributed to the design of the study, interpreted the sequencing data and substantially revised the manuscript. AG conceived the study, contributed to its design, interpreted the sequencing data, wrote part of the Background and Results sections and substantially revised the manuscript. All authors read and approved the final manuscript.

## Acknowledgements

We thank Diego Montiel Gonzalez for discussions and feedback. We thank the Princess Máxima Center for Pediatric Oncology, the patient and her parents for providing samples that made this research possible. We also appreciate the support of the Princess Máxima Center Biobank.

