## Additional File 2 for "Somatic variant detection using an individual-specific pangenome reference: a case study of a pediatric acute myeloid leukemia patient"

### 7_13 TRA breakends (hg38)

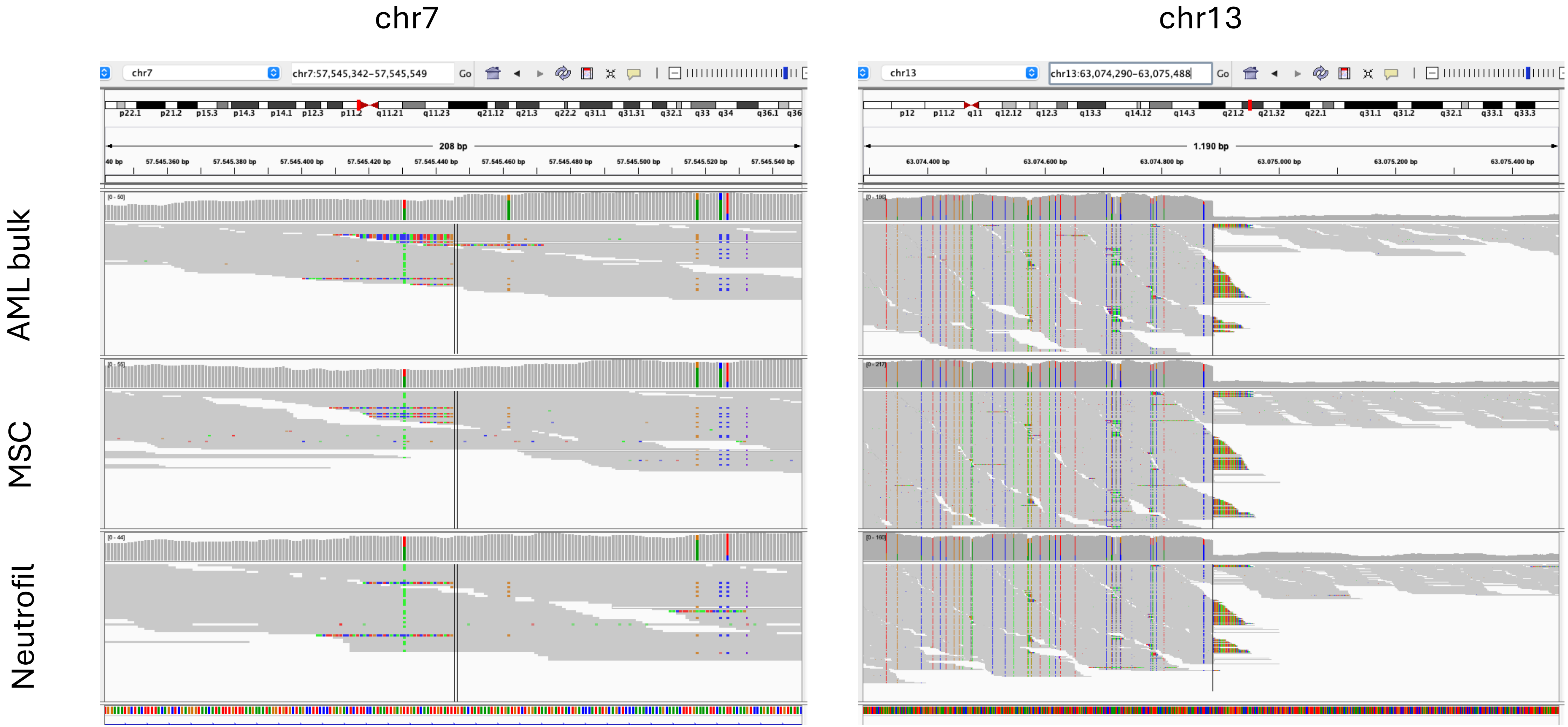

### 7_13 TRA breakends (pangenome)

chr7 chr13

AML bulk

MSC

Neutrofil

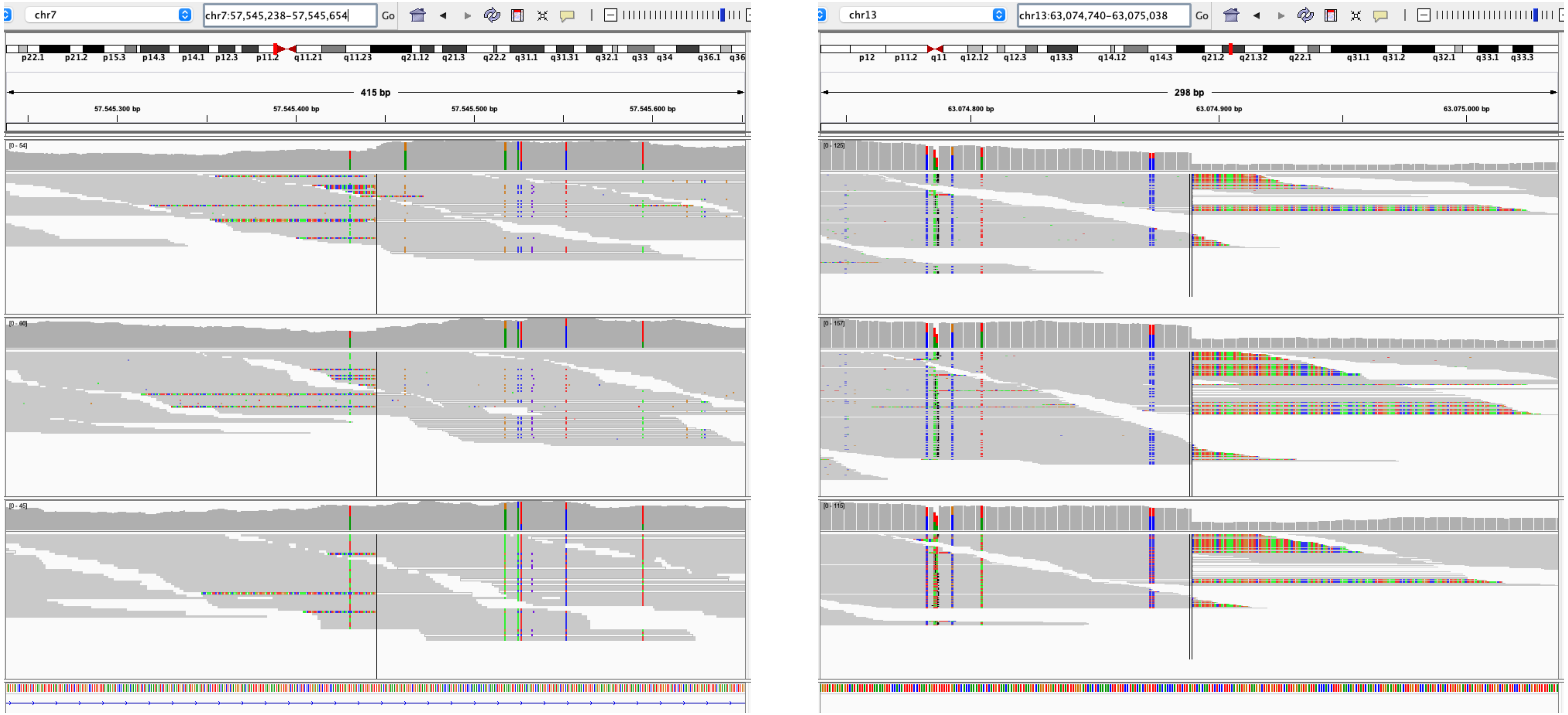

### 5_11_TRA breakends (hg38)

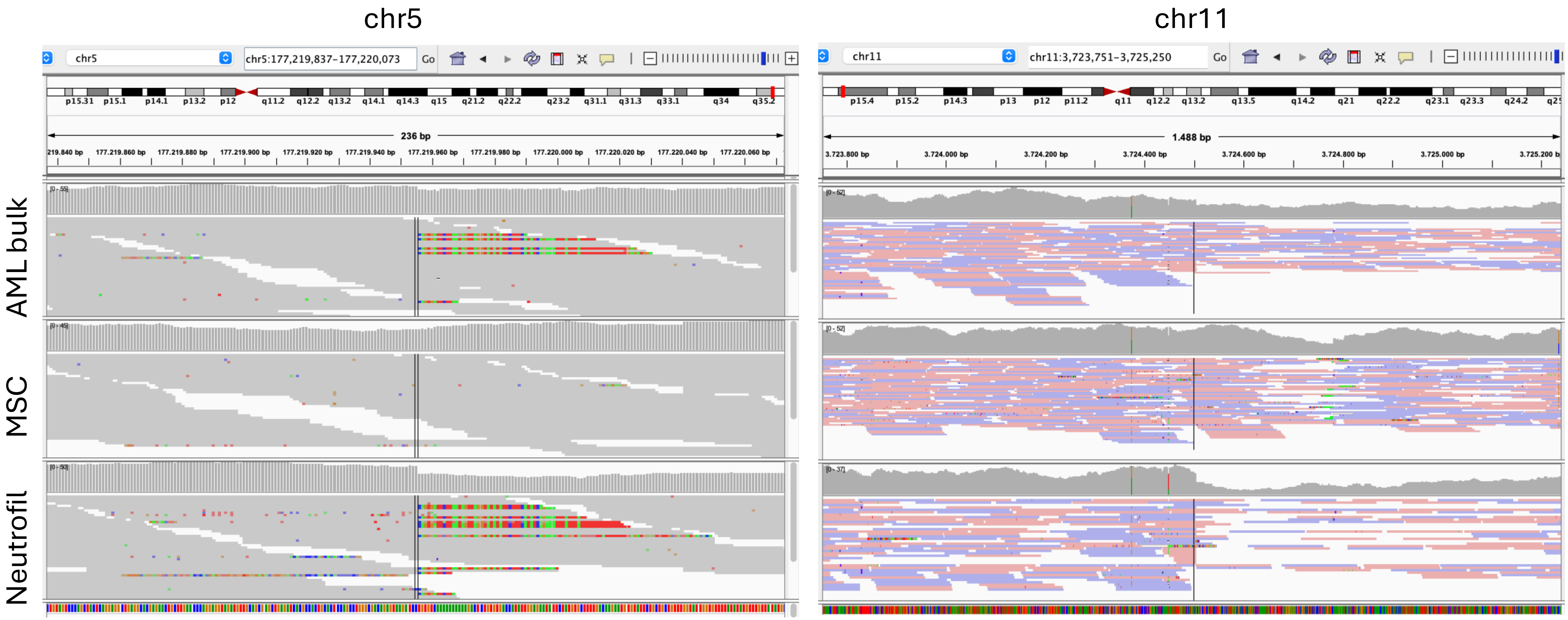

### 5_11_TRA breakends (pangenome)

chr5 chr11

AML bulk

MSC

Neutrofil

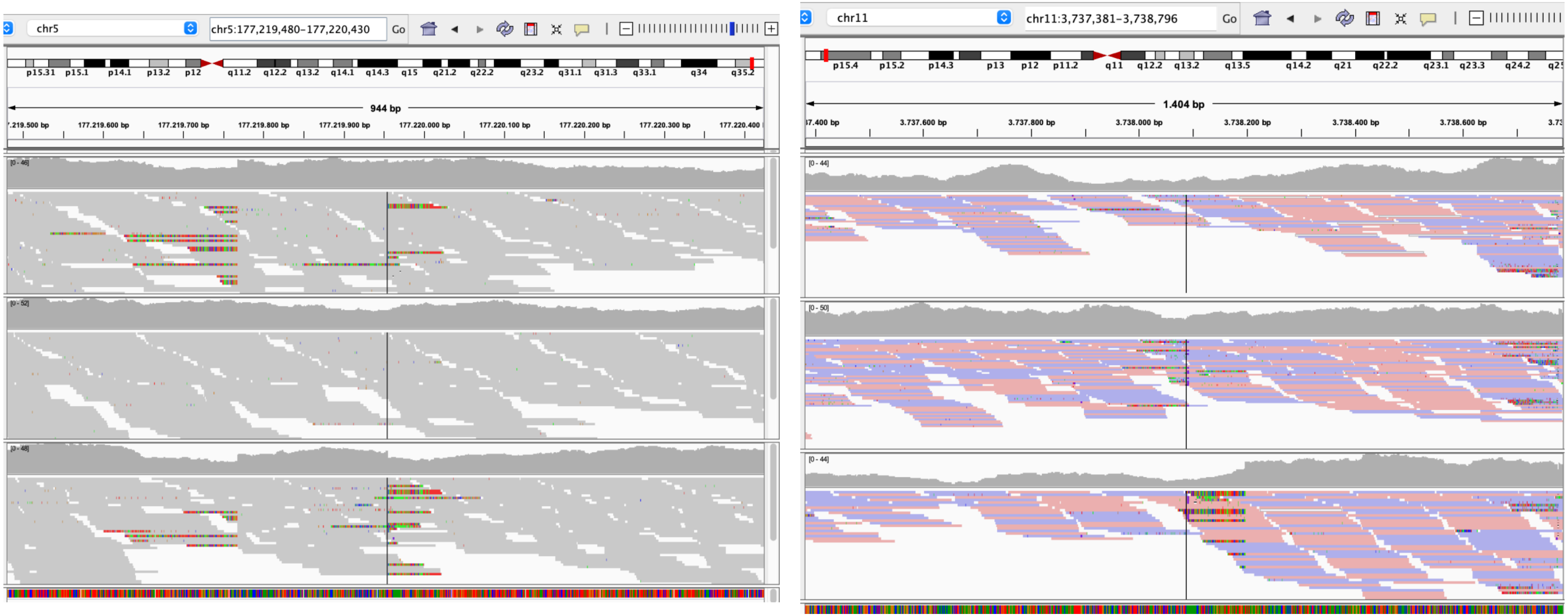

### 13_DUP breakends (hg38)

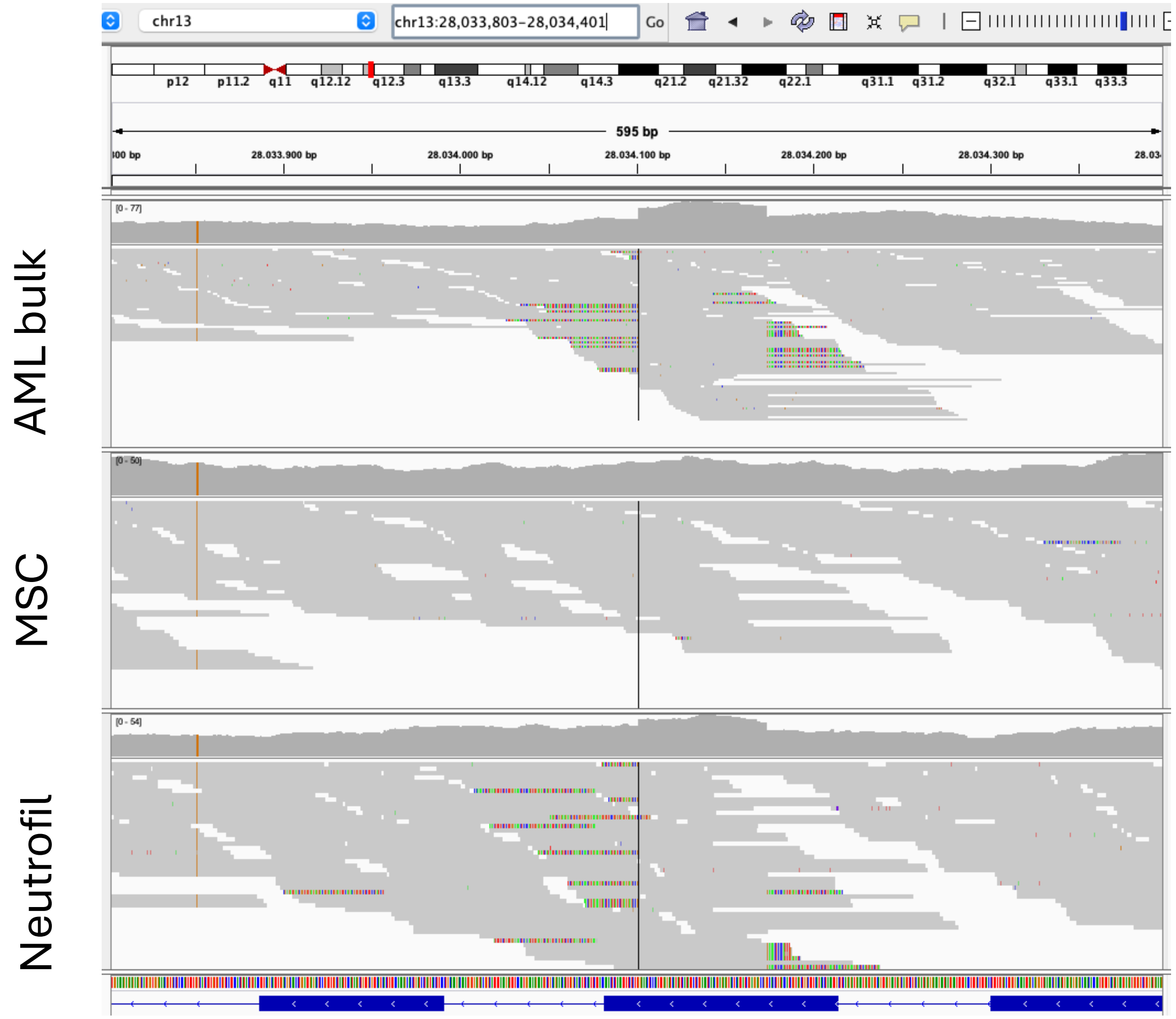

### 13_DUP breakends (pangenome)

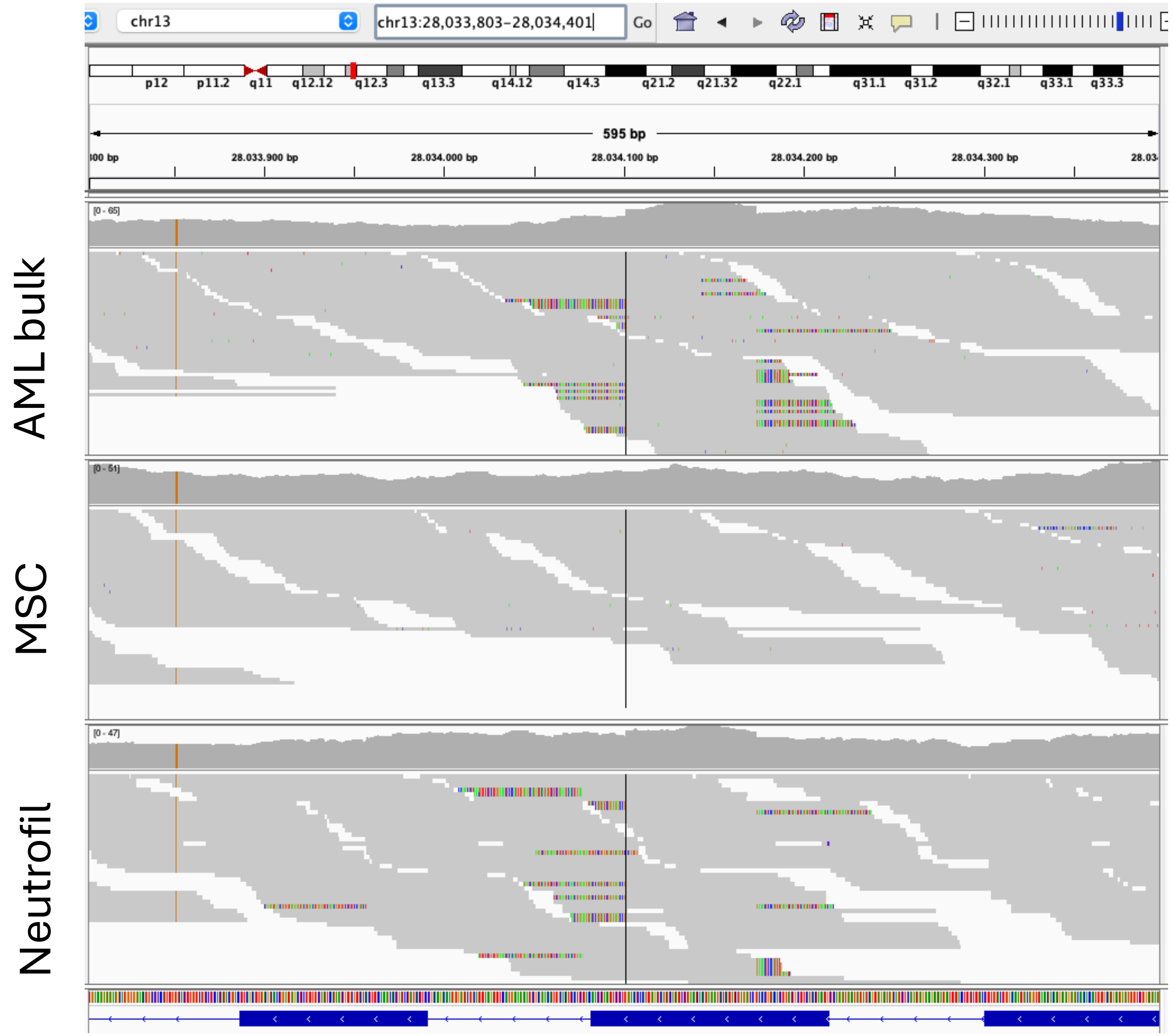

### 7_DEL breakends (hg38)

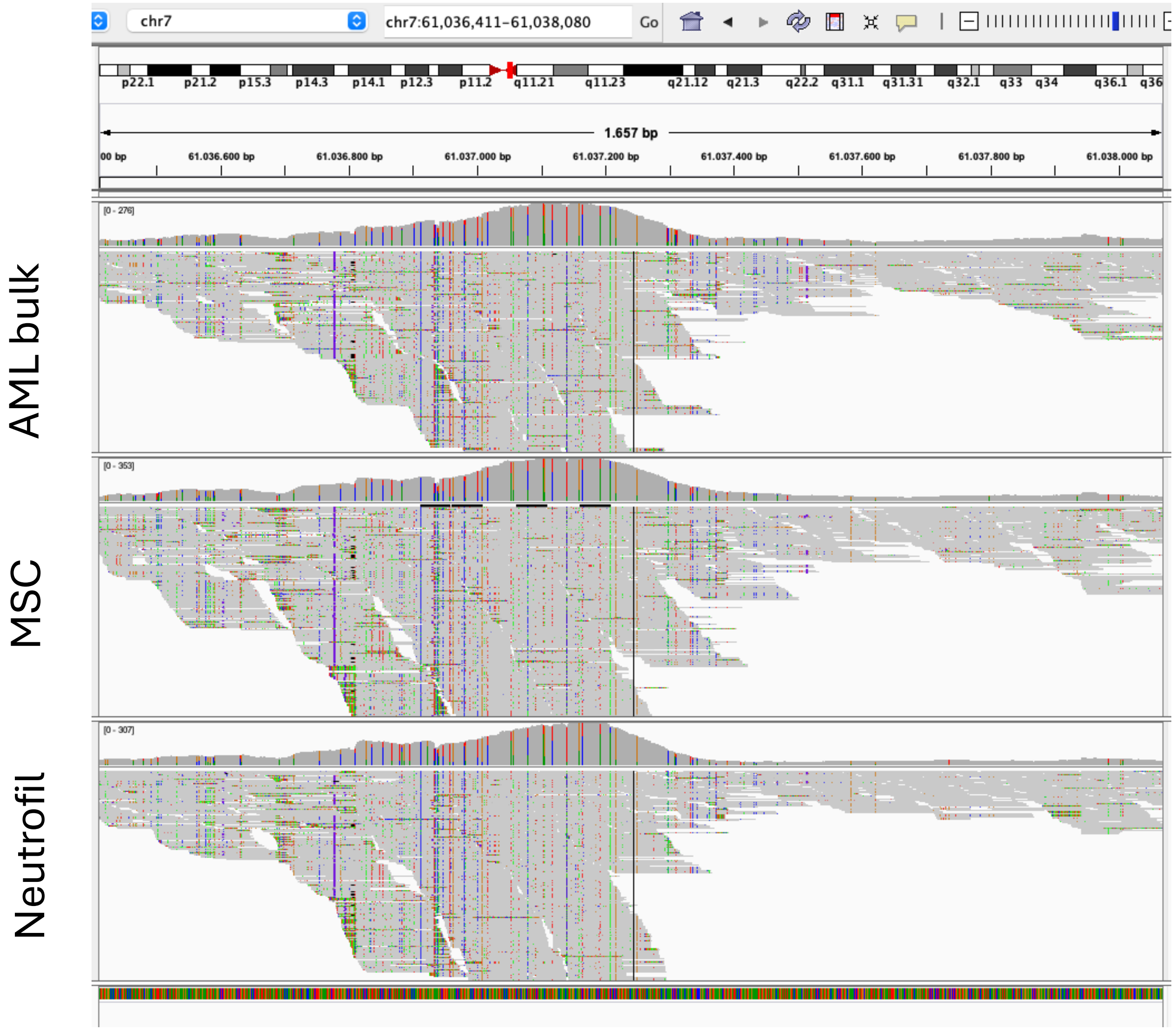

### 7_DEL breakends (pangenome)

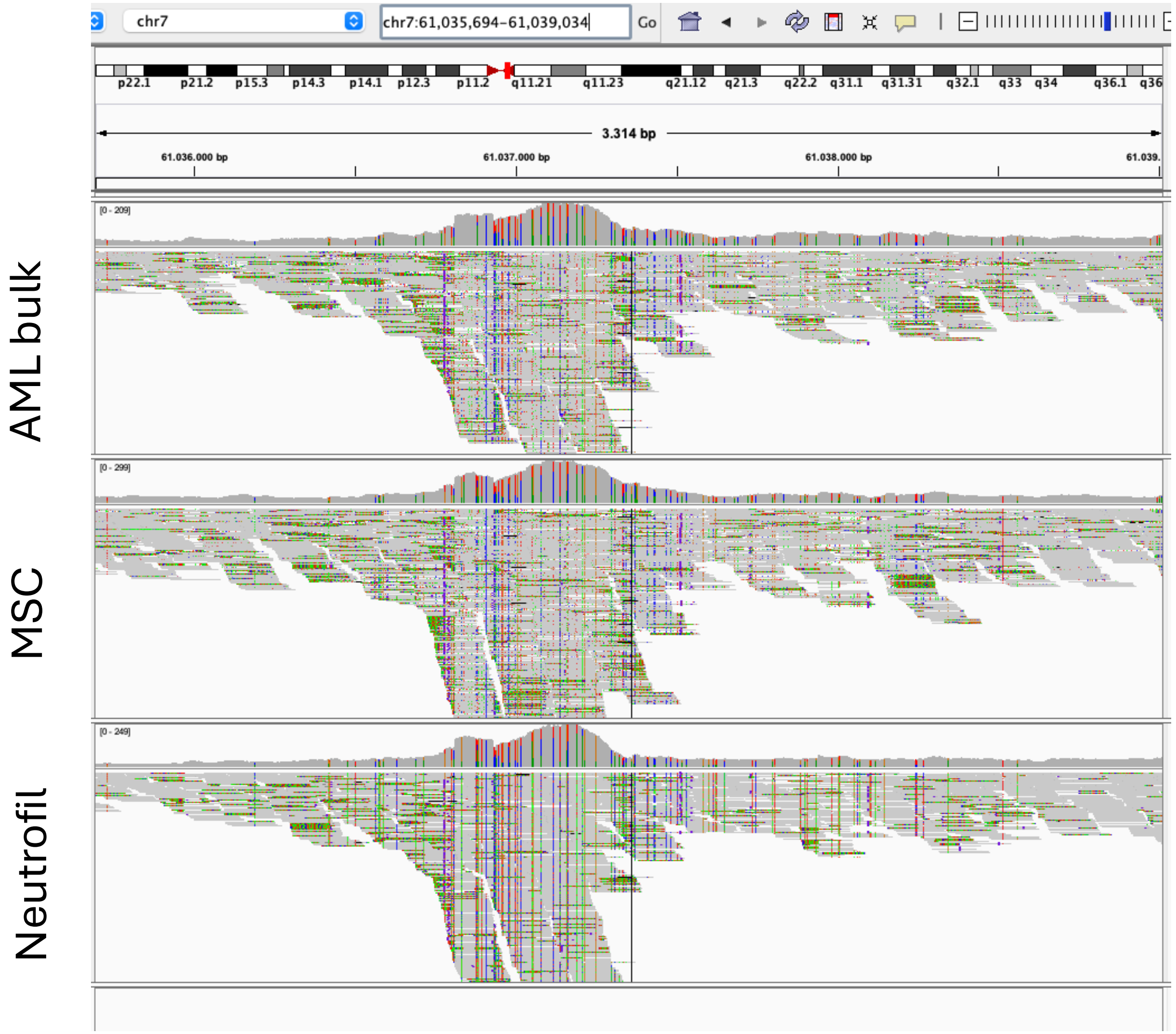

### 15_20_TRA breakends (hg38)

chr15

AML bulk

MSC

Neutrofil

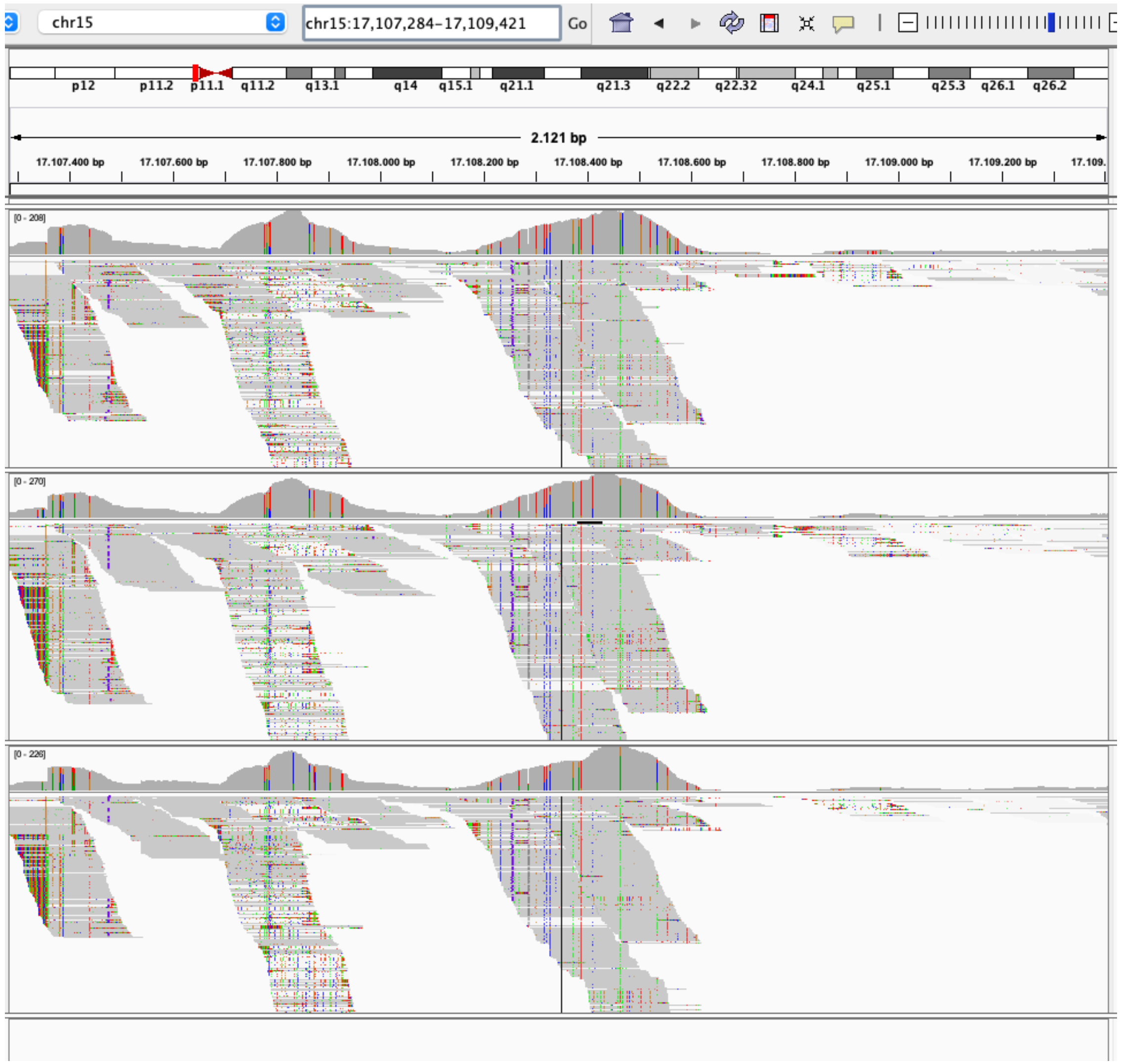


chr20

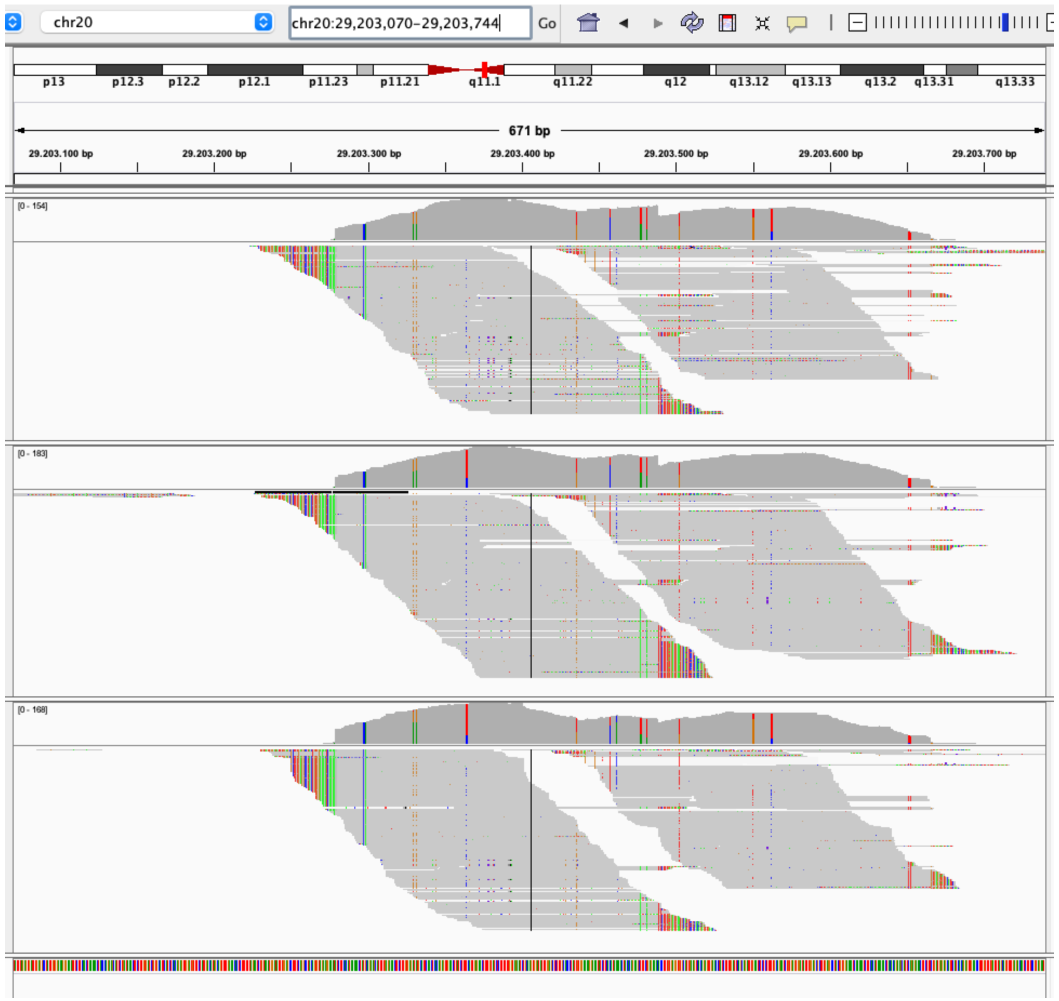

### 15_20_TRA breakends (pangenome)

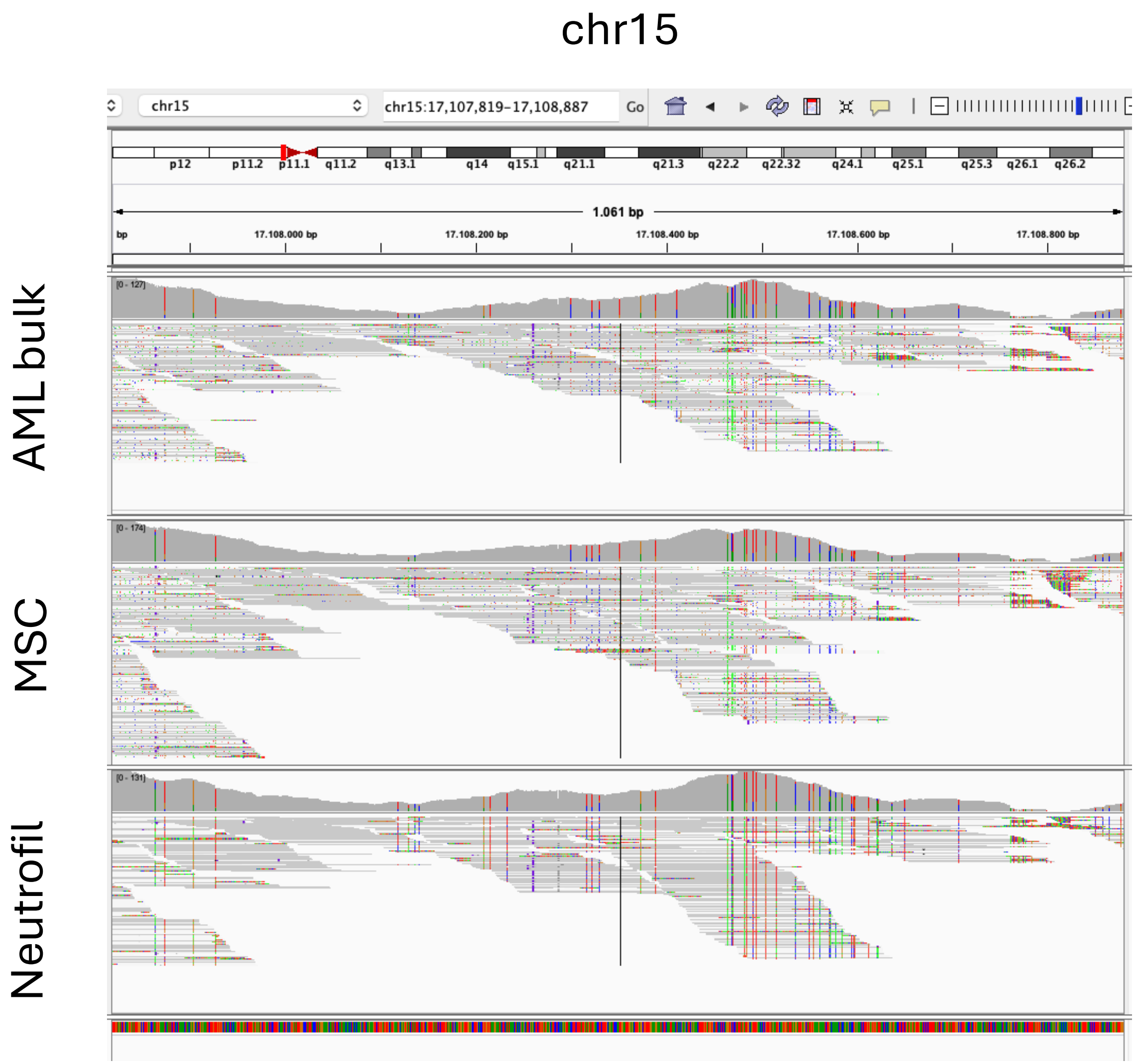


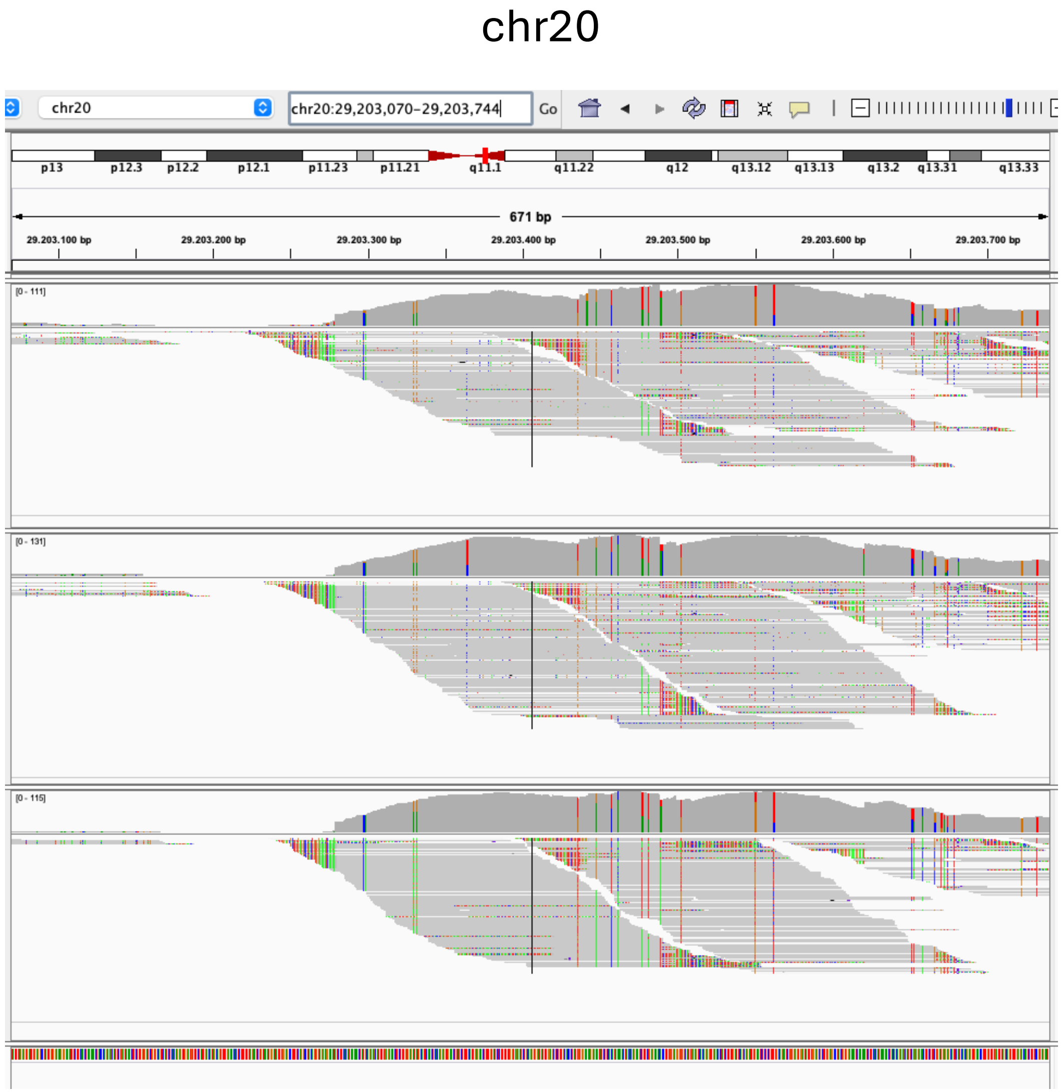

### 9_DEL/DUP breakends (hg38)

AML bulk

MSC

Neutrofil

SV start

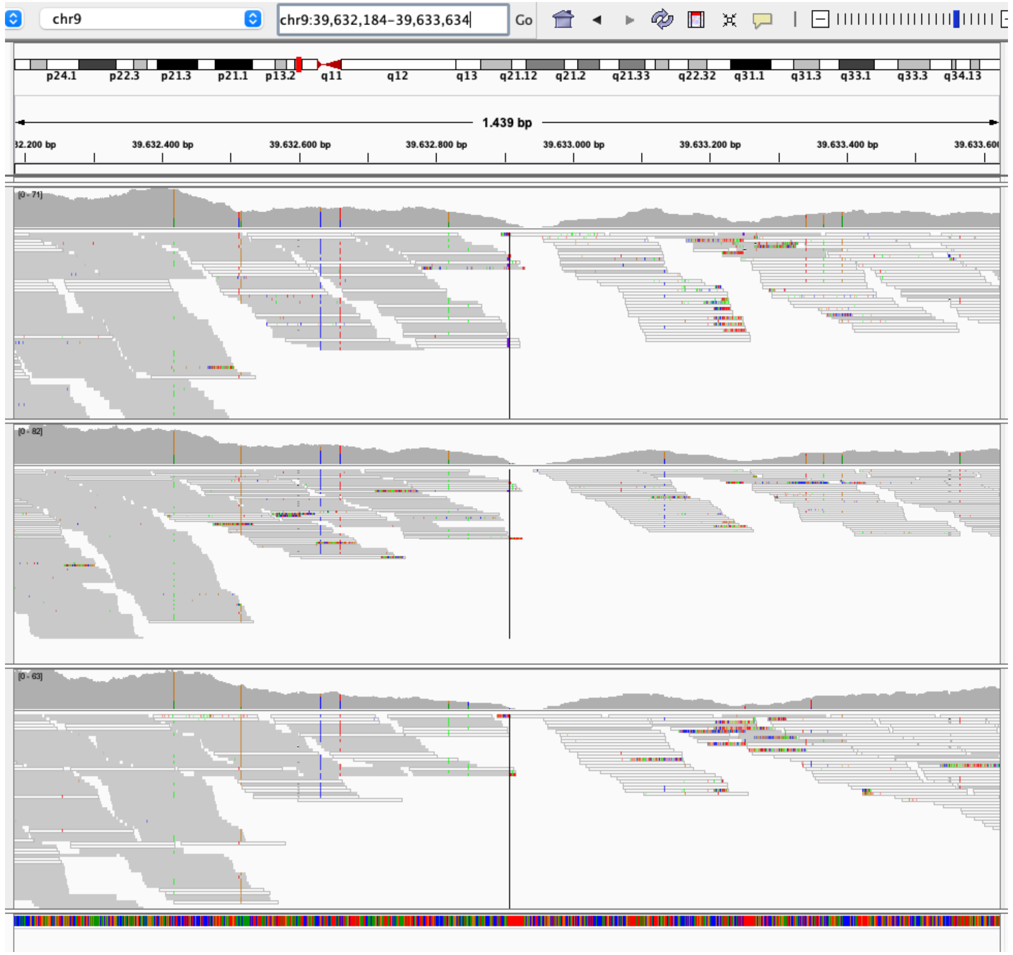


SV end

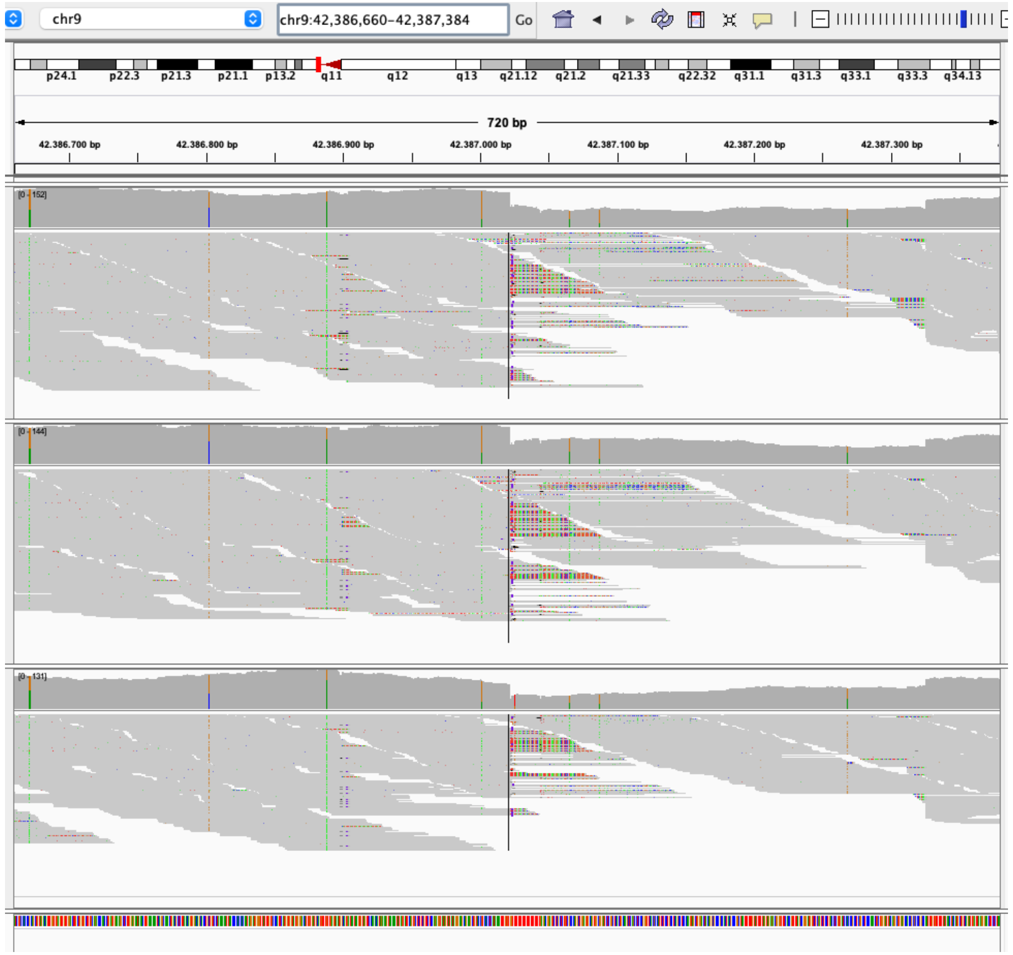

### 9_DEL/DUP breakends (pangenome)

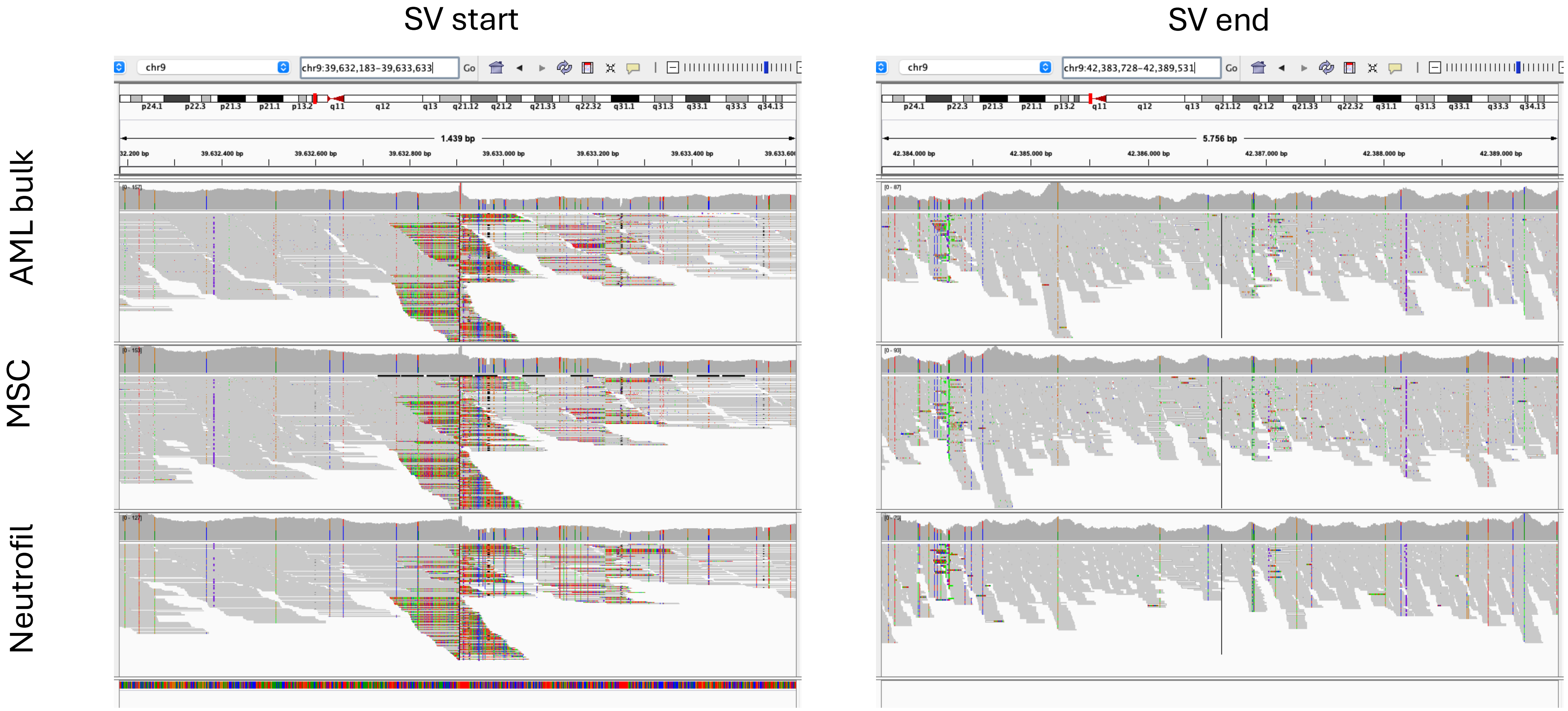

### 7_16_TRA breakends (hg38)

chr7

chr16

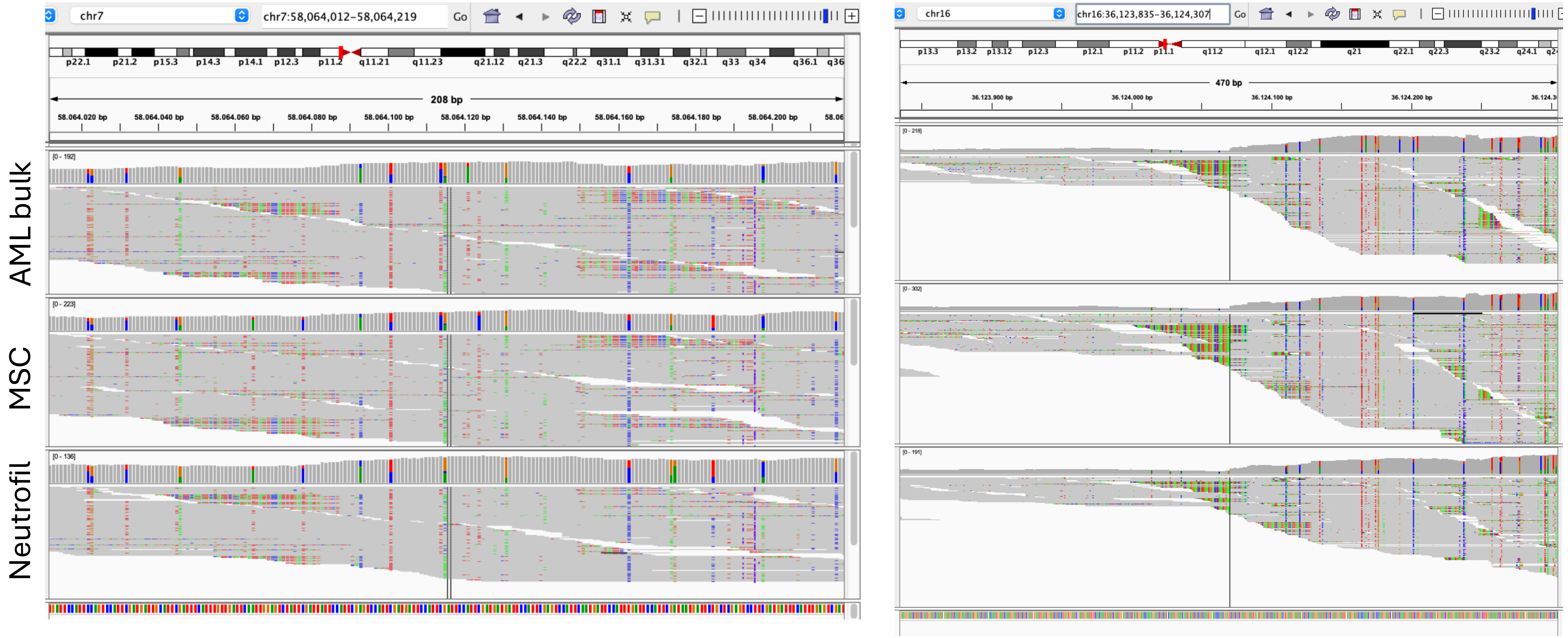

### 7_16_TRA breakends (pangenome)

chr7

chr16

AML bulk

MSC

Neutrofil

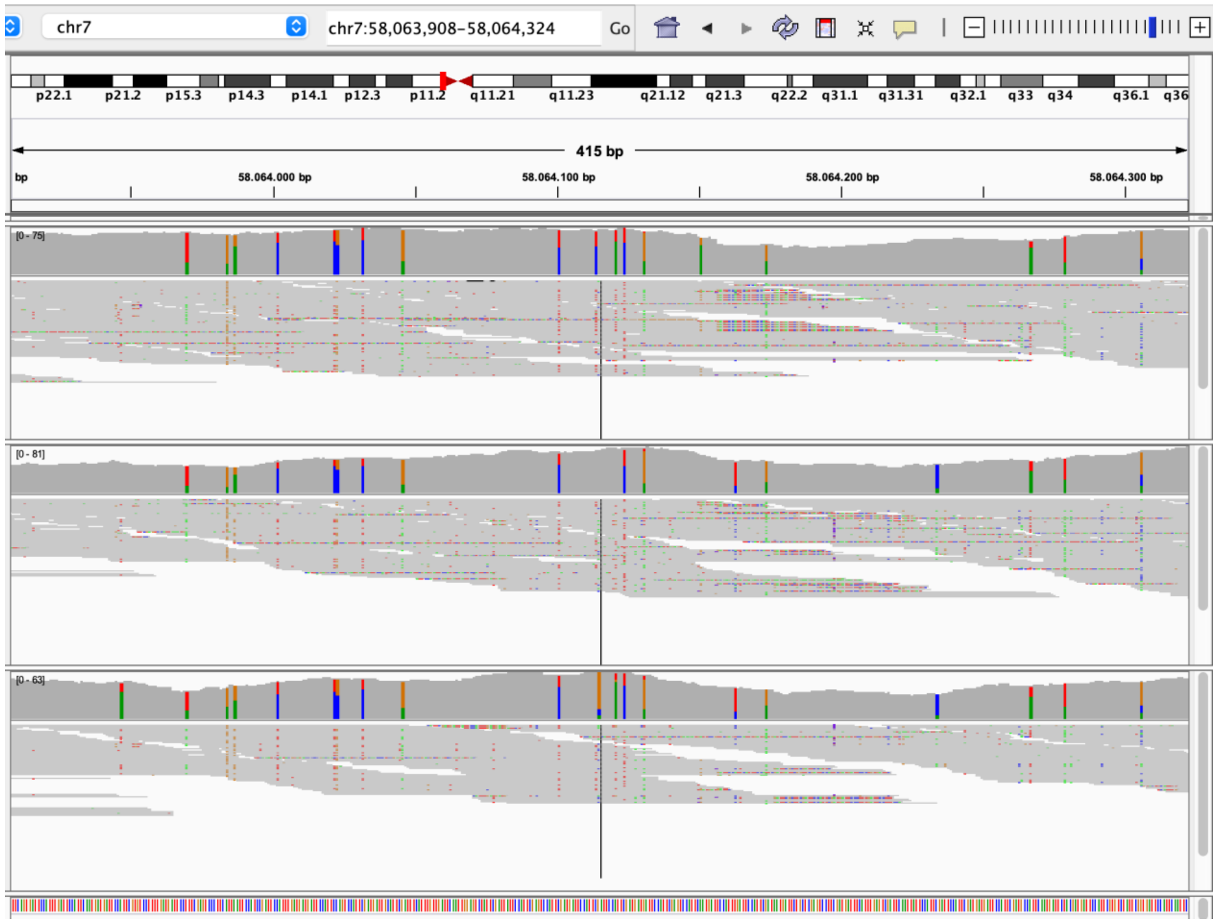


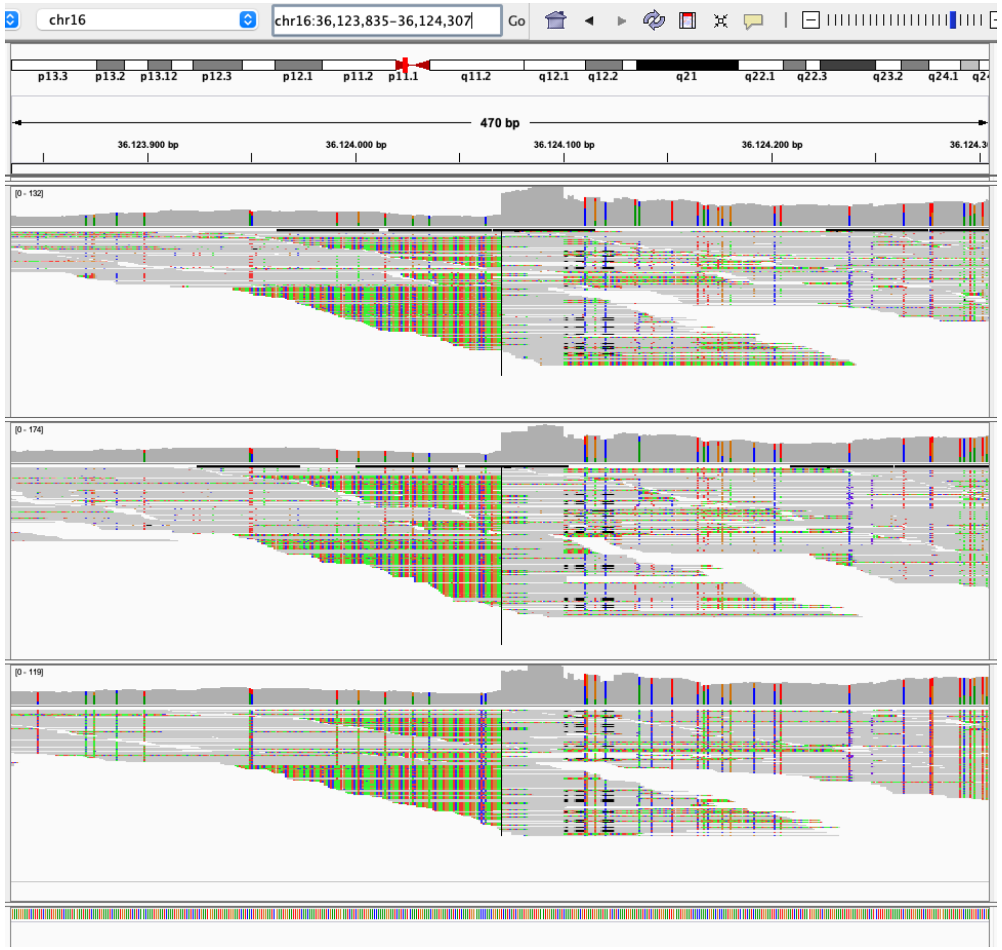

### 20_DEL breakends (hg38)

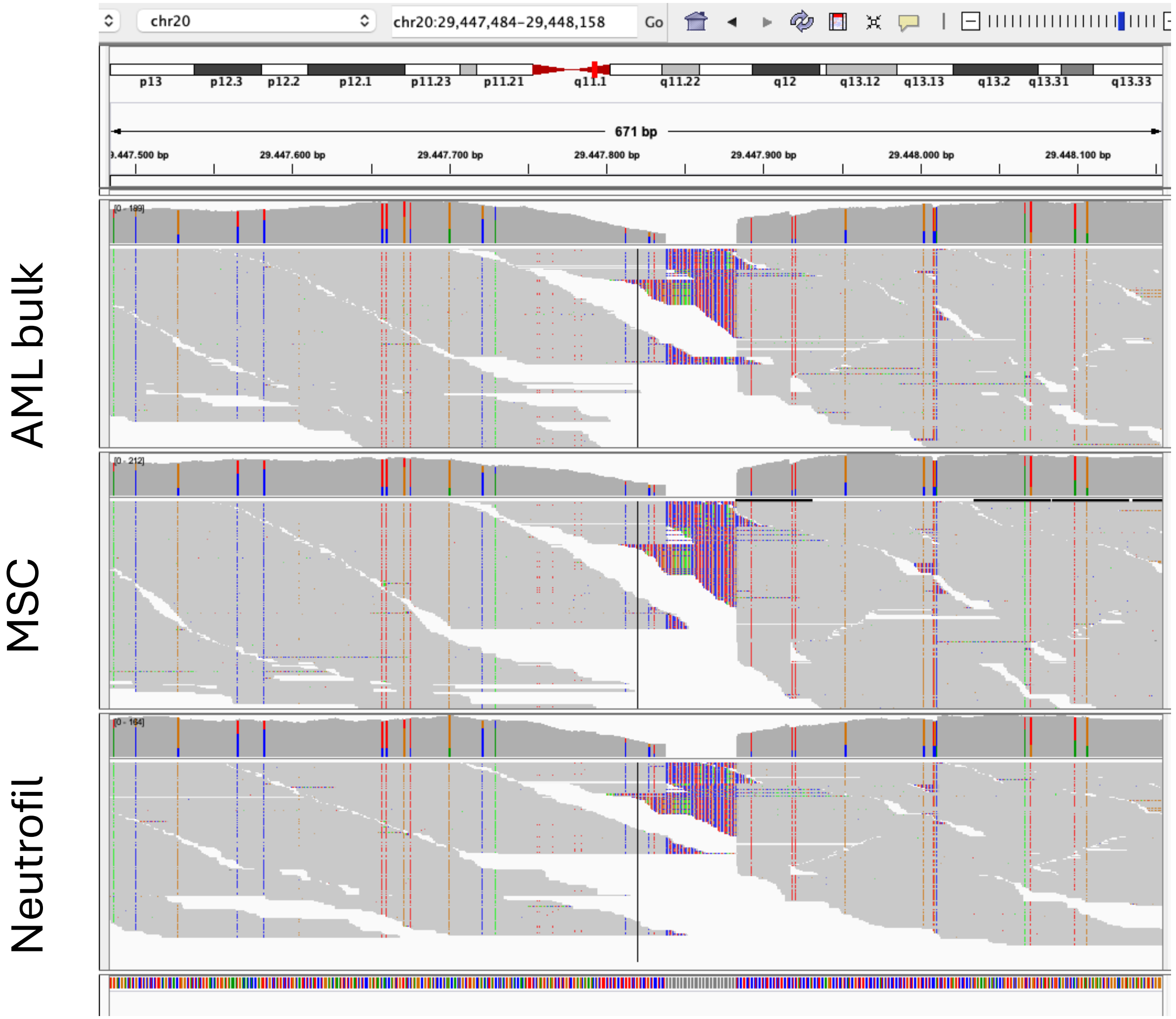

### 20_DEL breakends (pangenome)

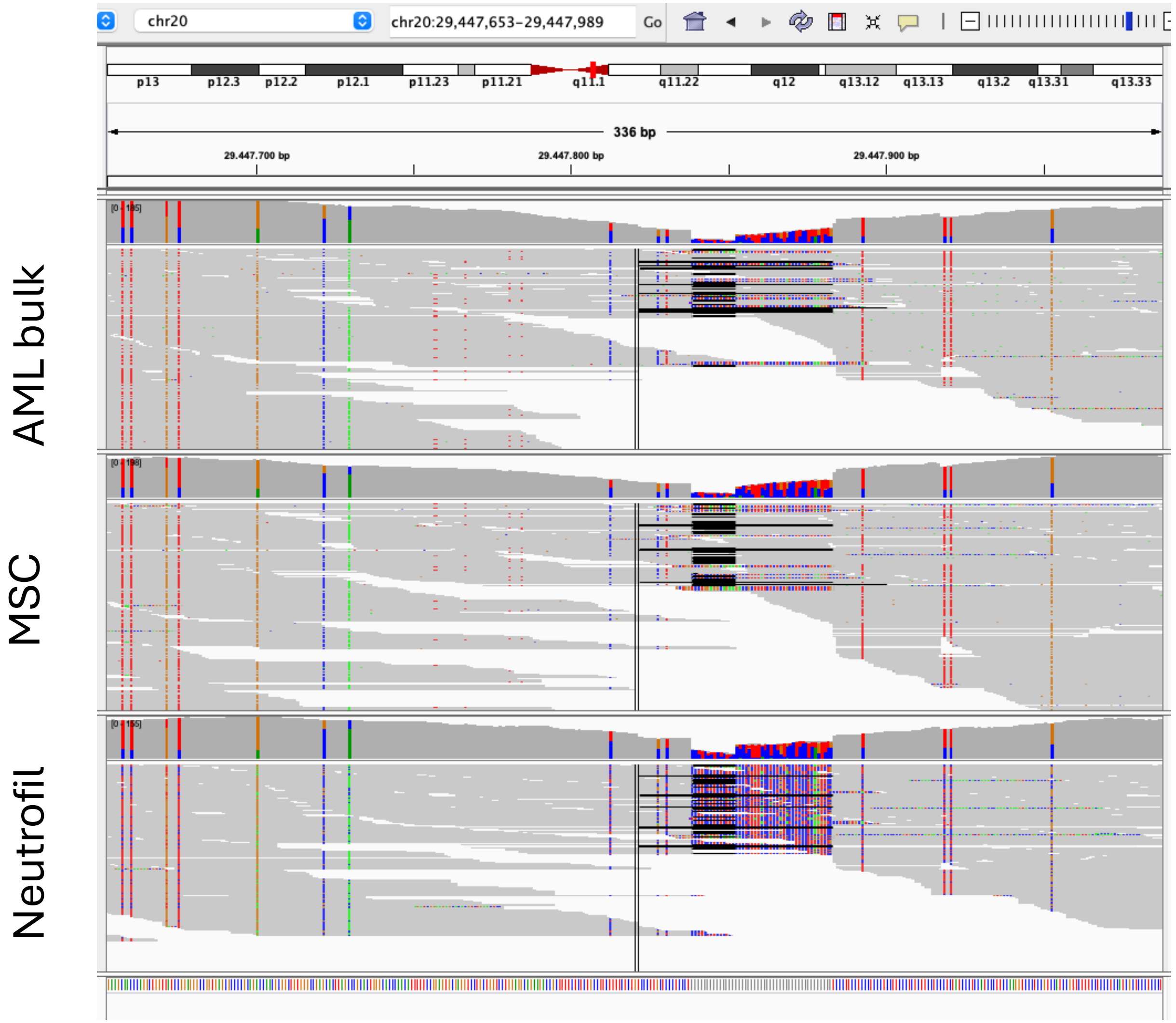
