## Additional File 1 for "Somatic variant detection using an individual-specific pangenome reference: a case study of a pediatric acute myeloid leukemia patient"

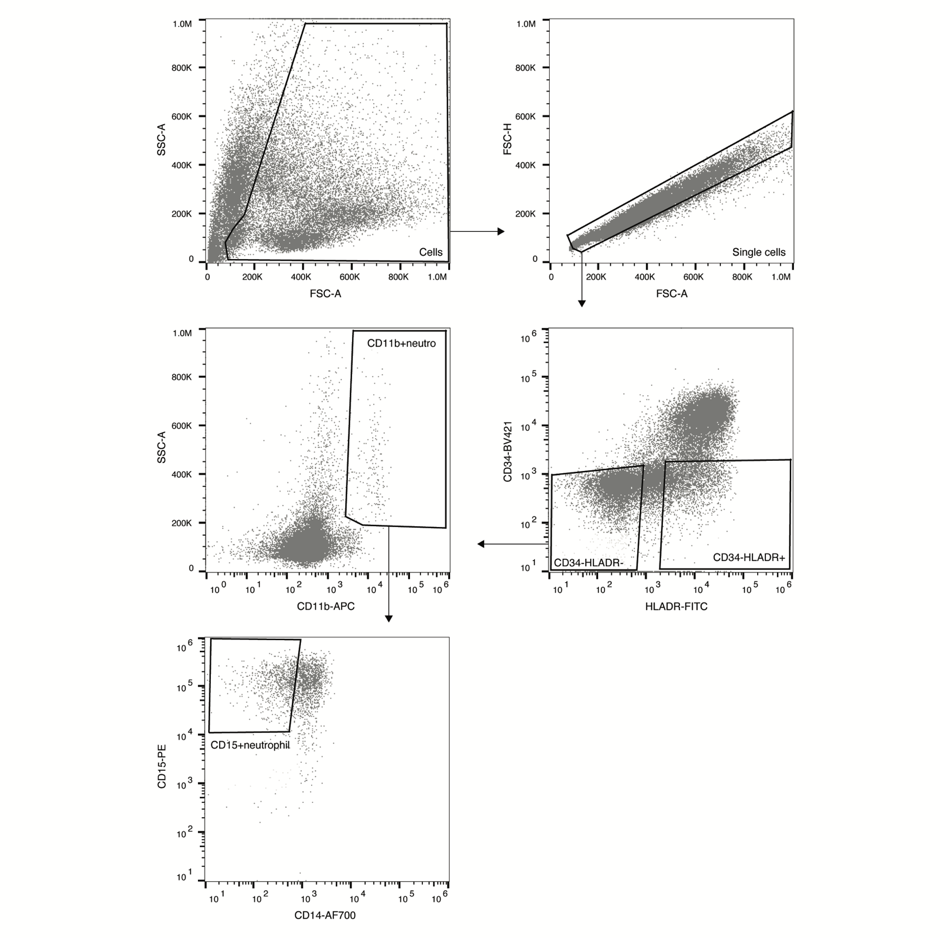

**Figure S1. Representative fluorescence activated cell sorting (FACS) plot for purification of neutrophils (CD34-HLADR-CD11b+CD14-).** Blasts in this patient sample were classified as (CD33+C38+CD45dimCD34+/-, CD117+, HLADR+/-).

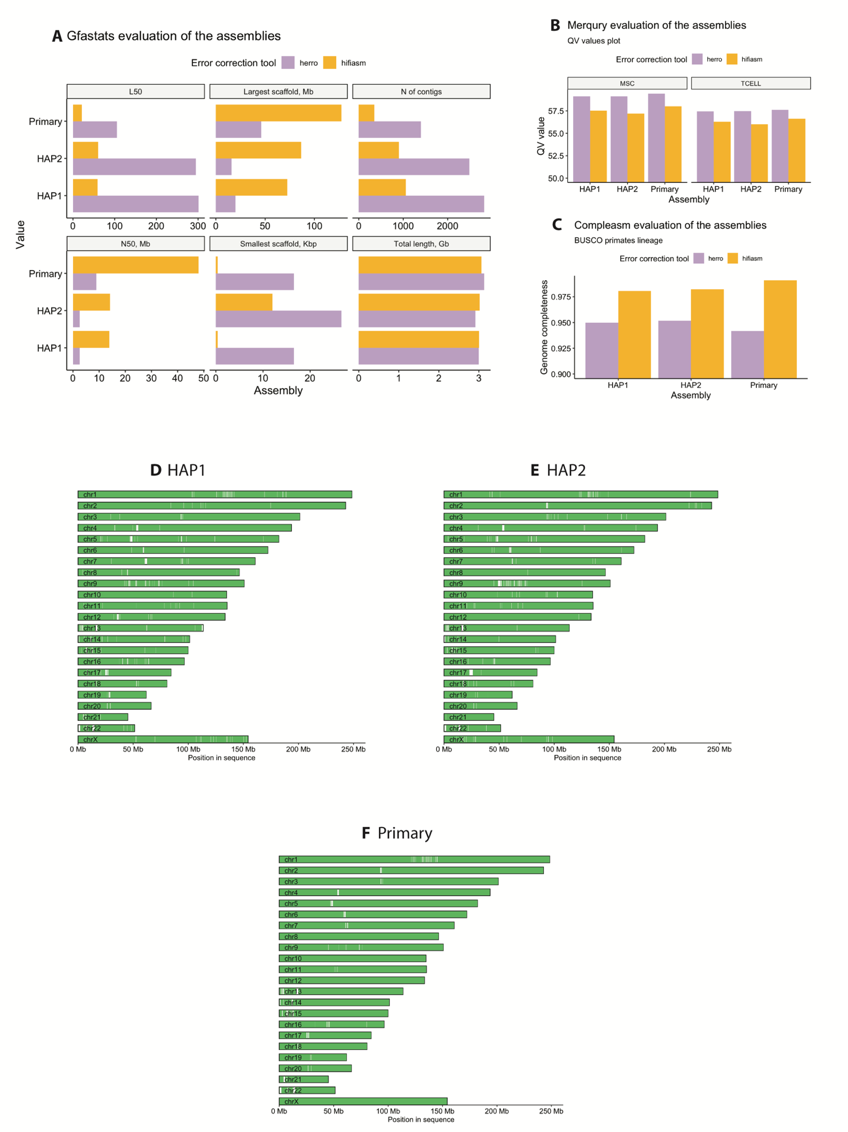

**Figure S2. Assemblies’ quality evaluation.** A – gfastats evaluation of haplotype-level (HAP1, HAP2) and primary assemblies (L50, N50, total number of contigs, lengths of the shortest and longest scaffolds, and the total length of the assembled genome). B – k-mer-based evaluation of the assemblies corrected with HERRO and hifiasm-ont with Merqury using short read data of two bulk samples (T-cell and MSC). C - Assemblies’ completeness estimation with the BUSCO primates lineage. D-F - pairwise alignment of the assemblies against T2T-CHM13v2.0 with minimap2.

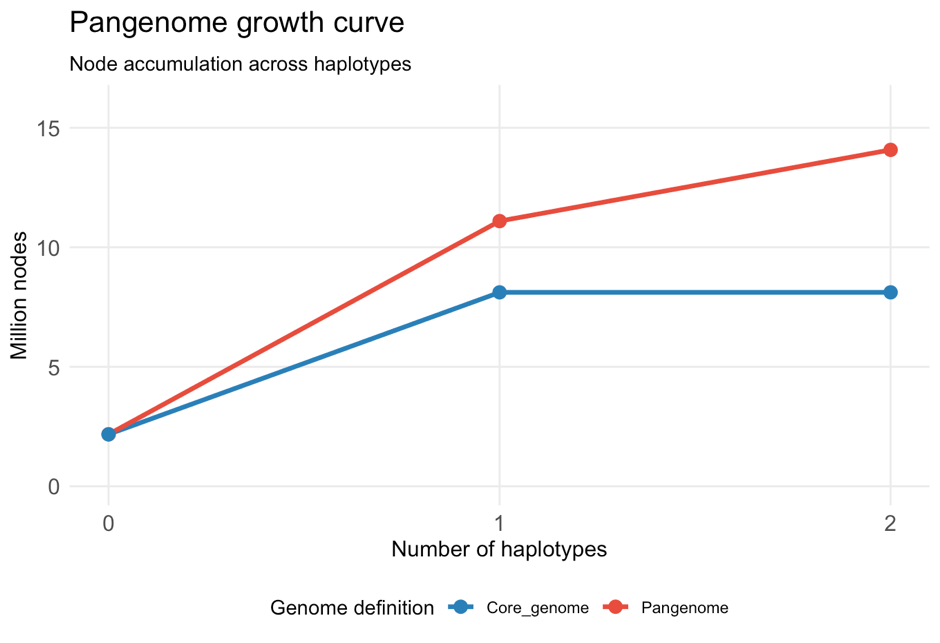

**Figure S3. Panacus growth curve for pangenome graph**. Curve showing the number of nodes added to the genome graph per each haplotype for hg38 (core genome, blue) and pangenome (red).

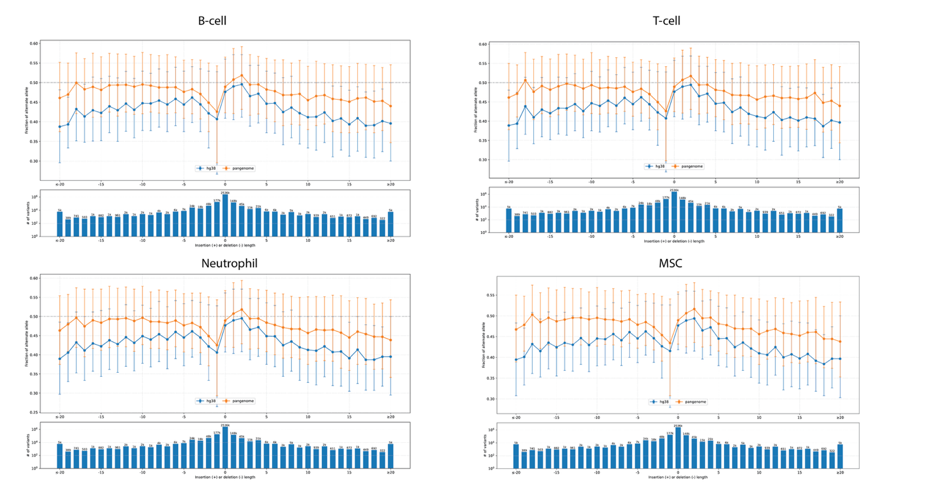

**Figure S4.** Bias-by-allele-length plot of hg38 (blue) and pangenome (orange) alignments for bulk samples: B-cells, T-cells, neutrophils and mesenchymal stromal cells.

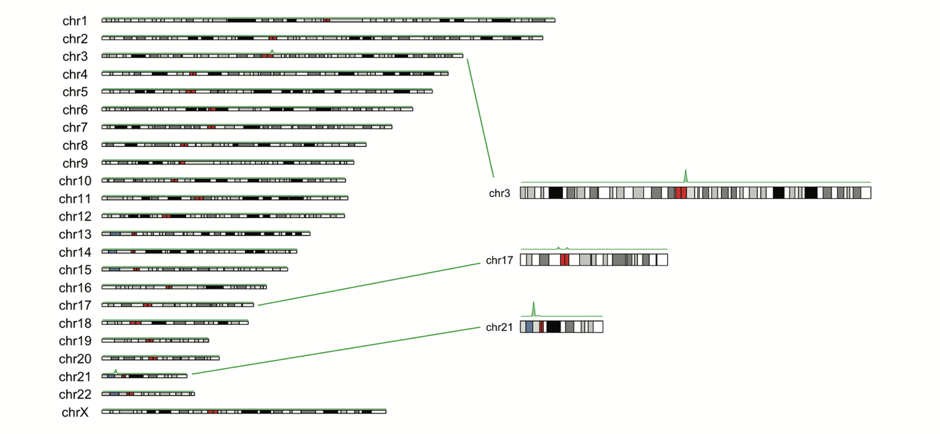

**Figure S5. Genomic location of rescued reads**. Ideogram of reads in five bulk samples rescued by mapping them on the pangenome graph and surjecting to hg38 coordinates. The green spikes indicate the enrichment of mapped surjected reads.

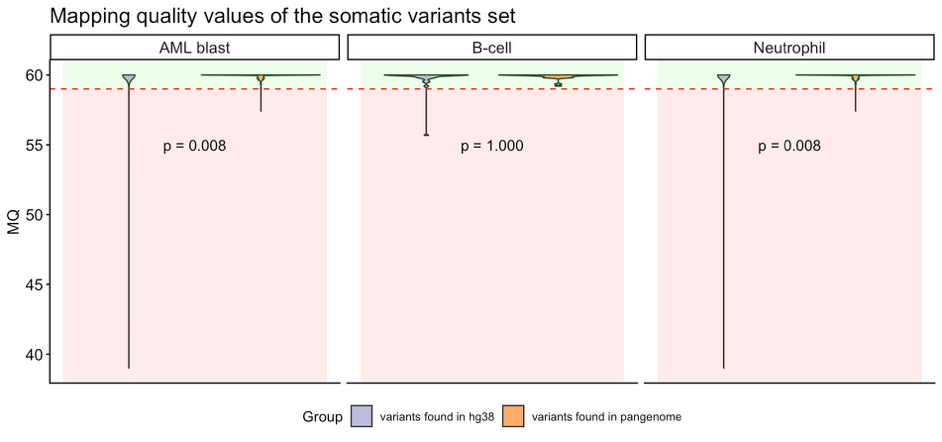

**Figure S6. Mapping quality (MQ) comparison for short variants called using linear and pangenome-based references (full plot).** The green zone indicates the variants that will be saved after applying a MQ cutoff = 59; the red zone indicates the variants that will be filtered out by the “BadMQ” tag.

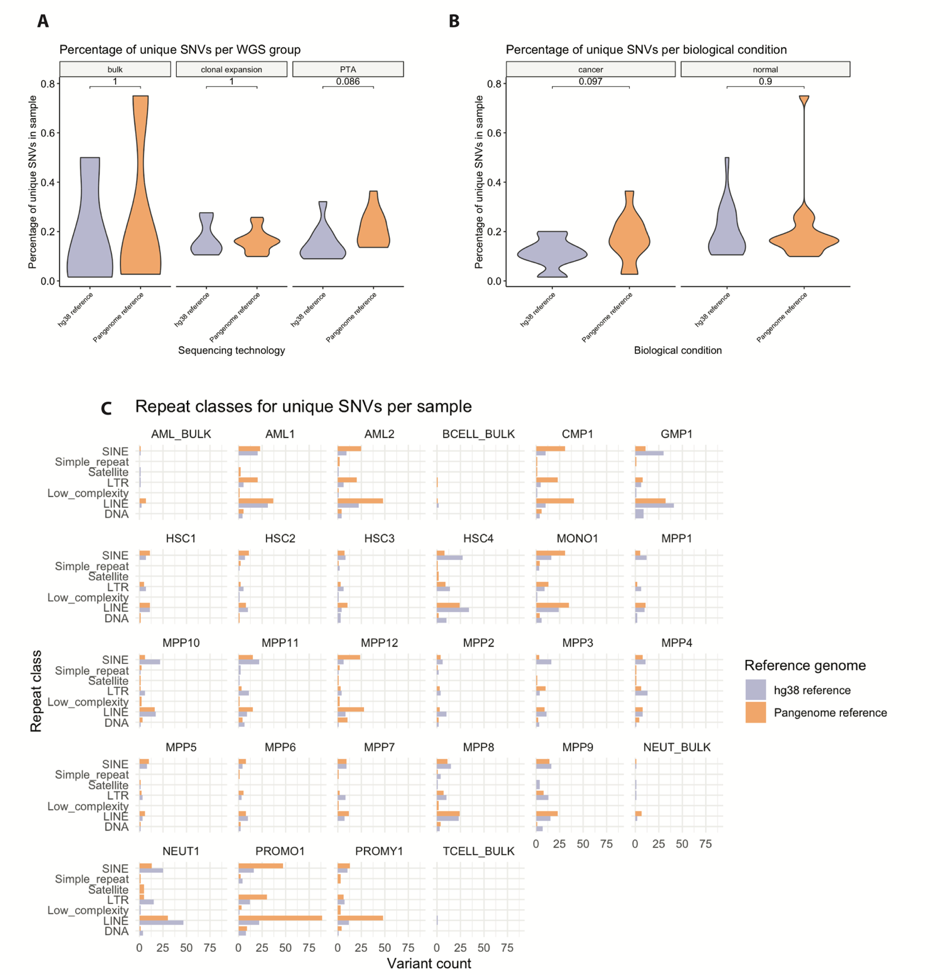

**Figure S7. Unique SNVs distribution.** Comparison of unique SNVs percentages per WGS sequencing group (A) and biological condition (B) between hg38-based and pangenome-based alignments (Mann-Whitney U test with Bonferroni correction). C – Absolute number of SNVs per sample across repeat classes in hg38-based and pangenome-based alignments.

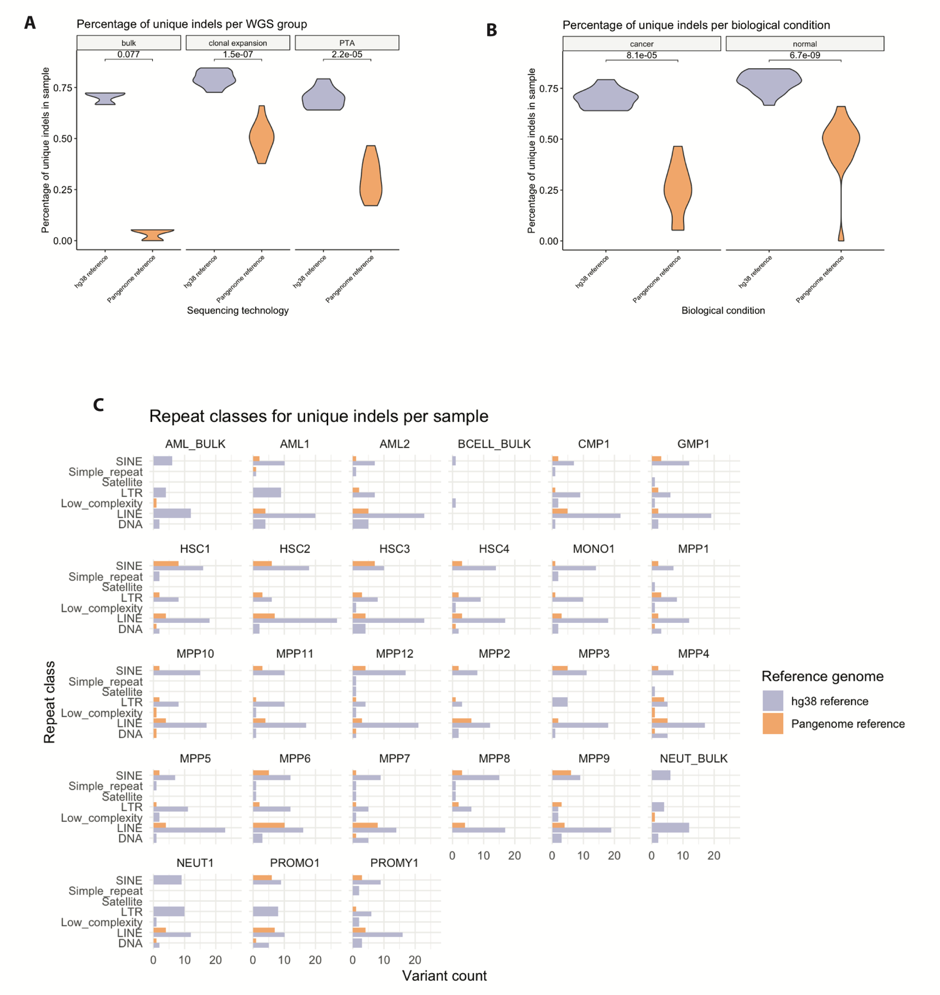

**Figure S8. Unique indels distribution.** Comparison of unique indels percentages per WGS sequencing group (A) and biological condition (B) between hg38-based and pangenome-based alignments (Mann-Whitney U test with Bonferroni correction). C – Absolute number of indels per sample across repeat classes in hg38-based and pangenome-based alignments.

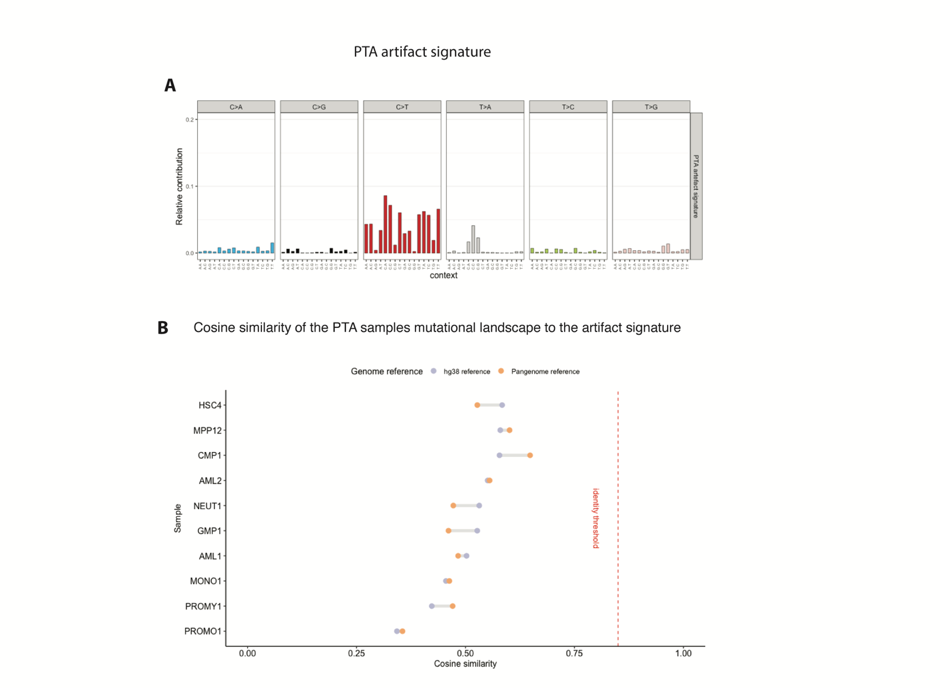

**Figure S9. Primary template amplification (PTA) artifact assessment.** A – SBS mutational signature of the PTA artifact introduced by strand displacement polymerases. B – Comparison of cosine similarity values to the artifact signature for PTA-derived samples aligned using the hg38 and pangenome references. The red dashed line shows the cosine similarity identity threshold for the PTA signature.

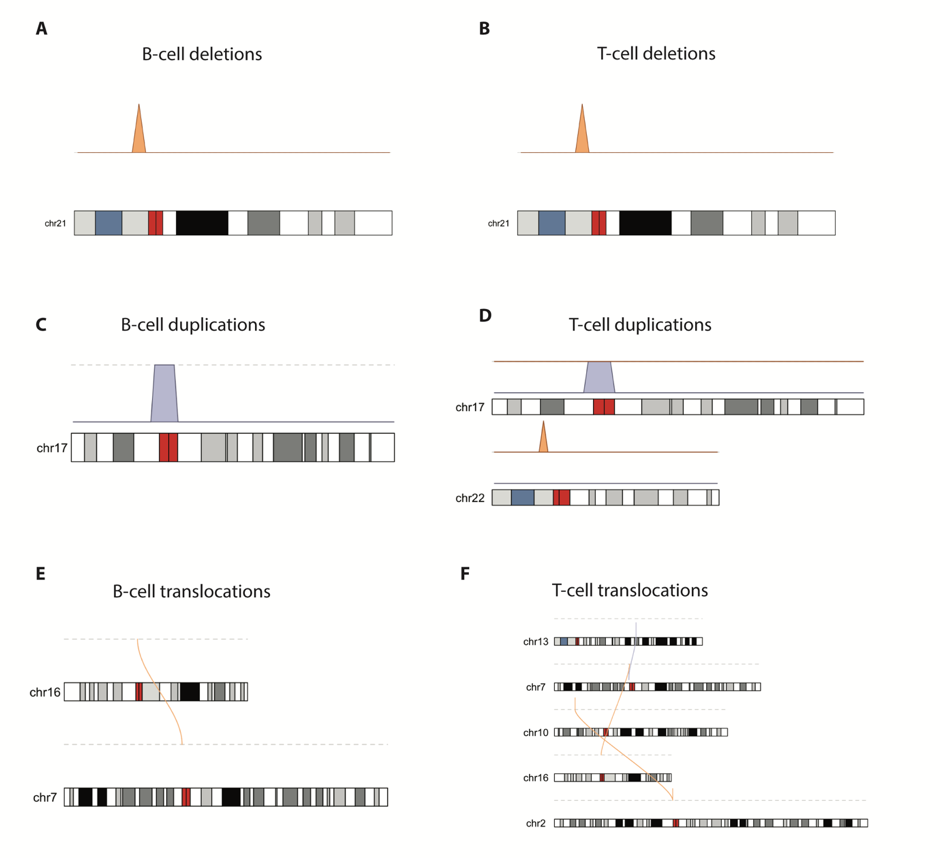

**Figure S10. Ideograms of somatic SV events in B-cell and T-cell samples**. SVs identified with pangenome-based and hg38-based analyses: deletions (A,B), duplications (C,D), and translocations (E,F). Driver SV events are shown in red.

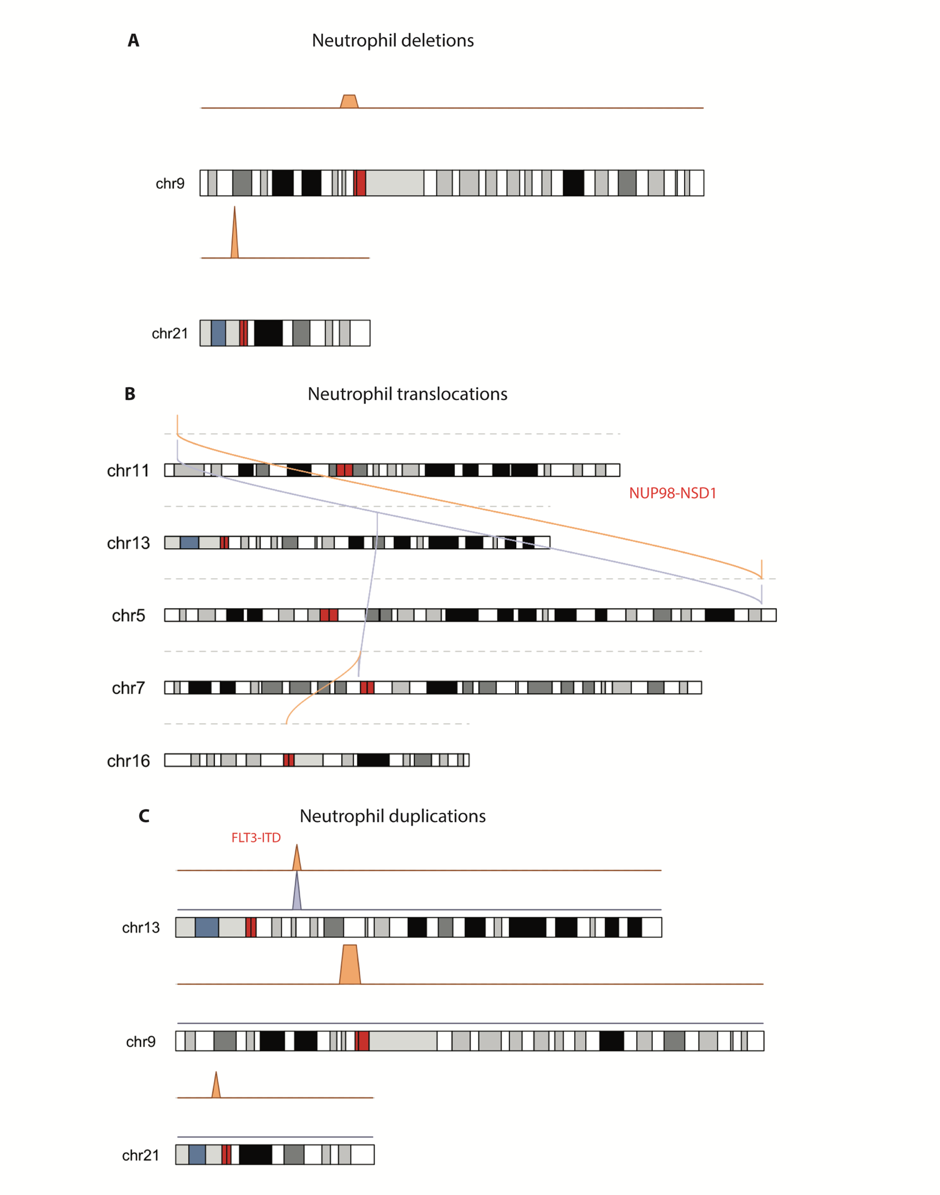

**Figure S11. Ideograms of somatic SV events in neutrophil bulk sample identified with pangenome-based and hg38-based analyses.** Events shown include deletions (A), translocations (B), and duplications (C). Driver SVs are shown in red.

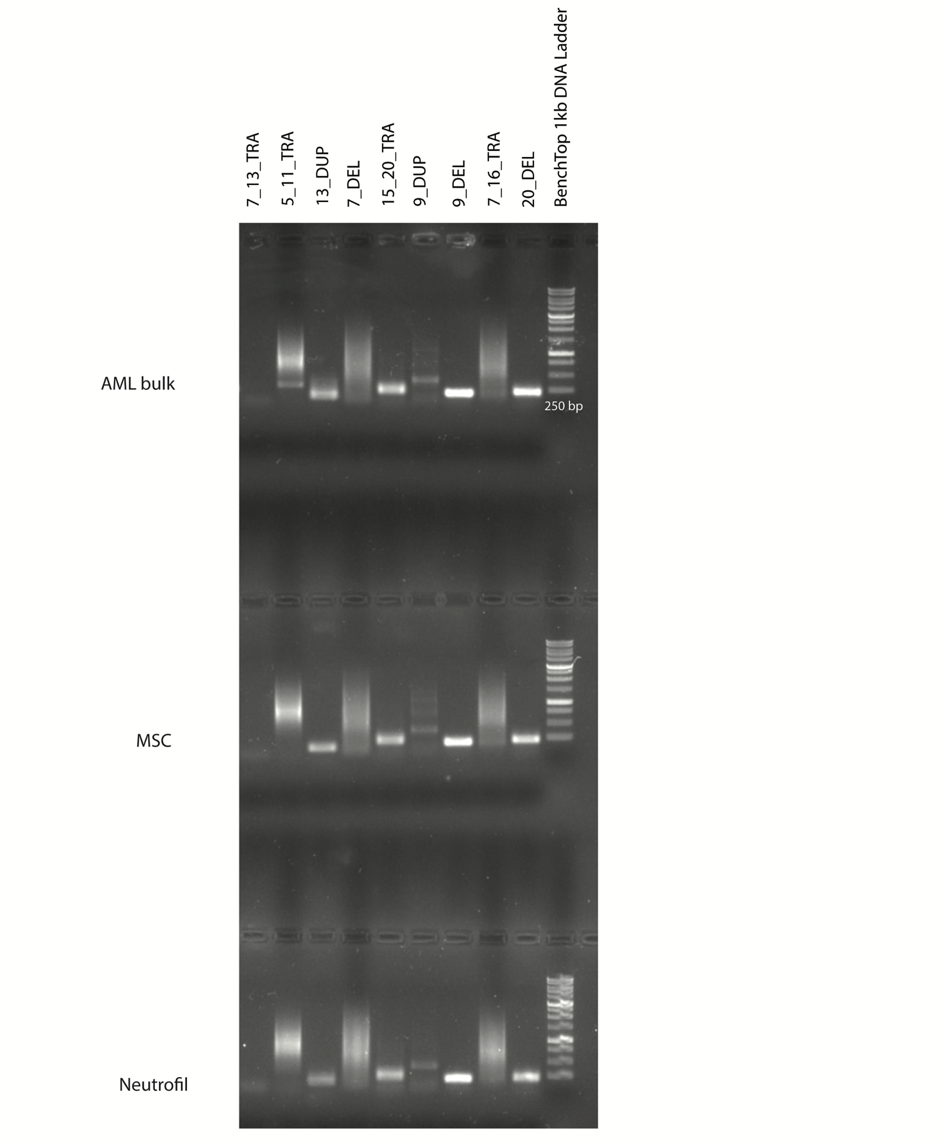

**Figure S12. PCR amplification bands for the junction fragments in AML bulk, neutrophil bulk and MSC bulk samples.** Each lane corresponds to one SV event: 5_11_TRA corresponds to the *NUP98-NDS1* translocation; 13_DUP corresponds to *FLT3-ITD*.

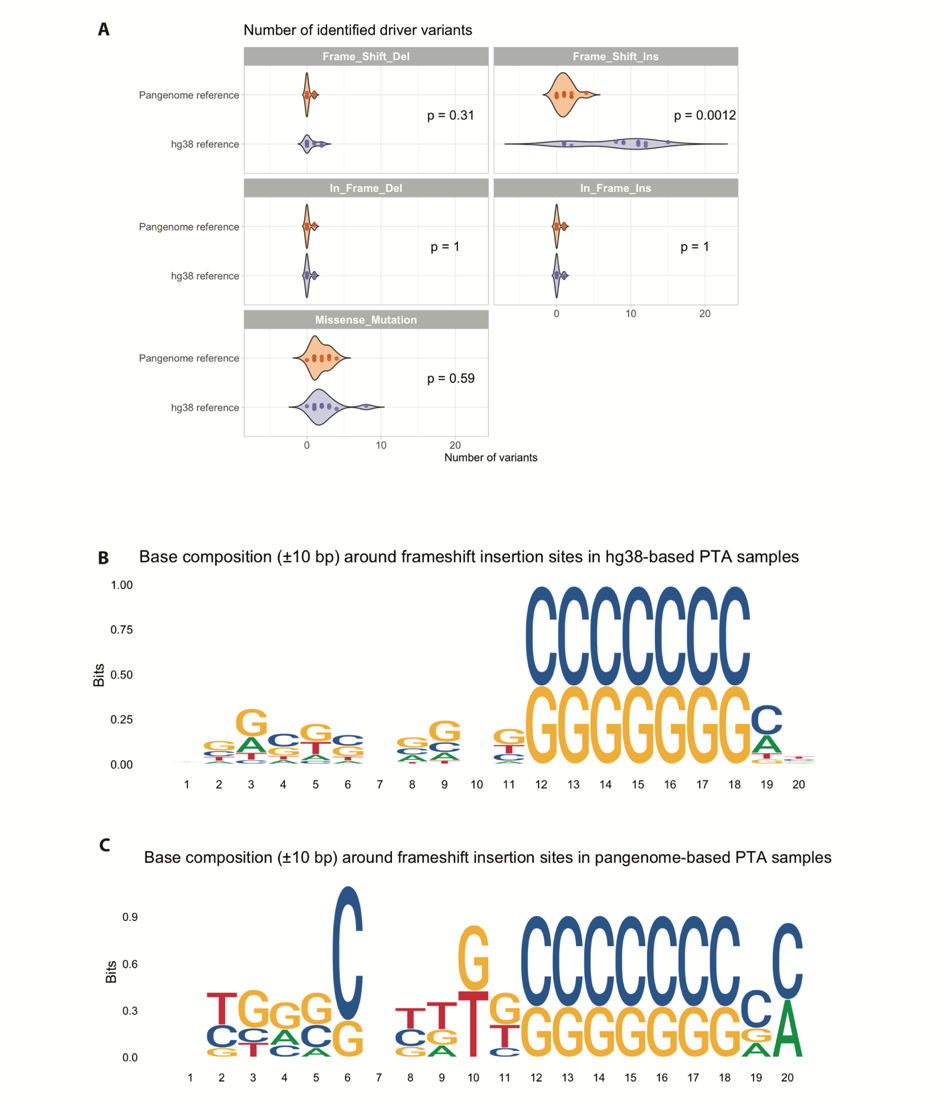

**Figure S13. Driver variants mutation type and context.** A – violin plots of driver variants distributions per mutation type (Mann-Whitney U test with Bonferroni correction). B – 10-bp sequence logo for frameshift insertions in hg38-based samples amplified with PTA. C - same sequence logo in pangenome-based samples amplified with PTA.

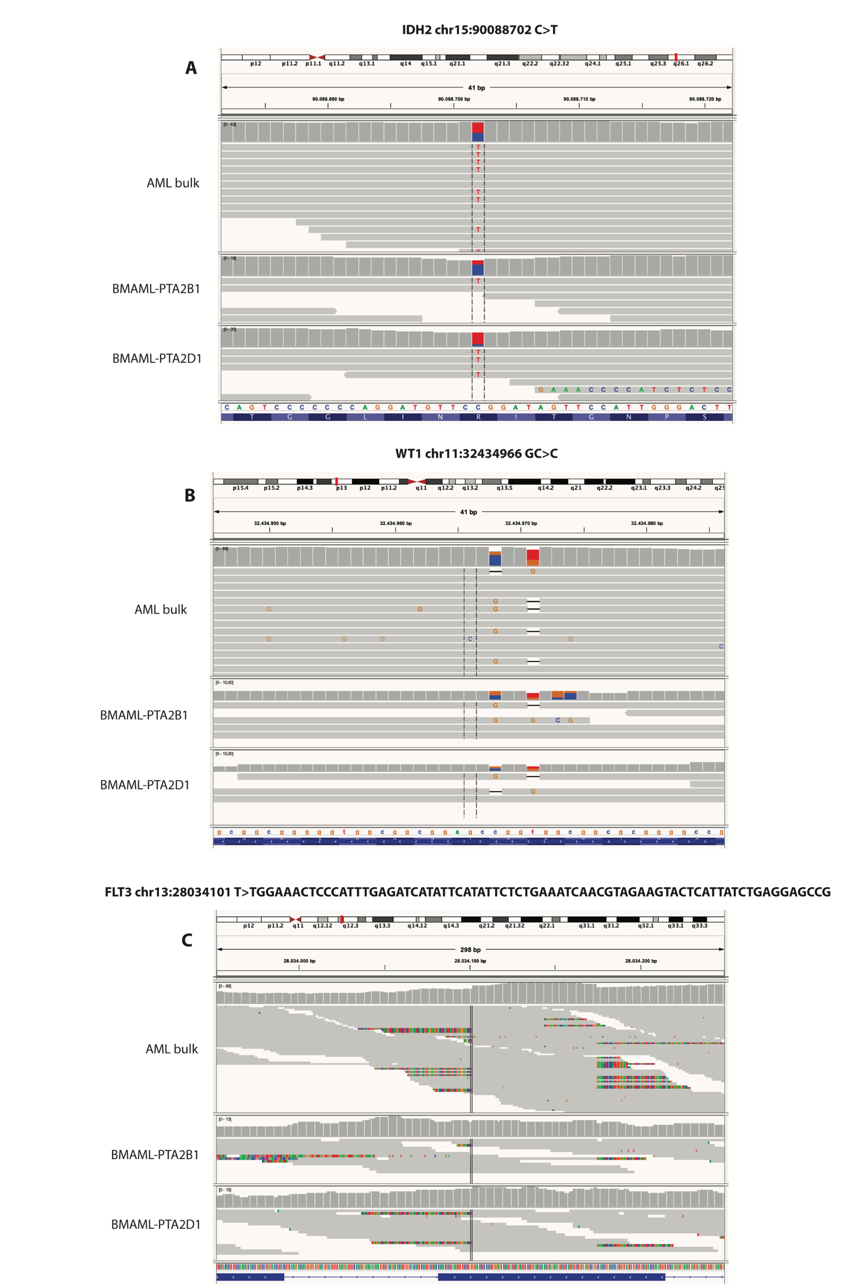

**Figure S14. Driver variants coverage in AML blasts.** Canonical driver variants in *IDH2* (A), *WT1* (B) and *FLT3* (C) genes in pangenome-based alignments of the AML bulk sample and single AML blasts samples amplified with PTA.

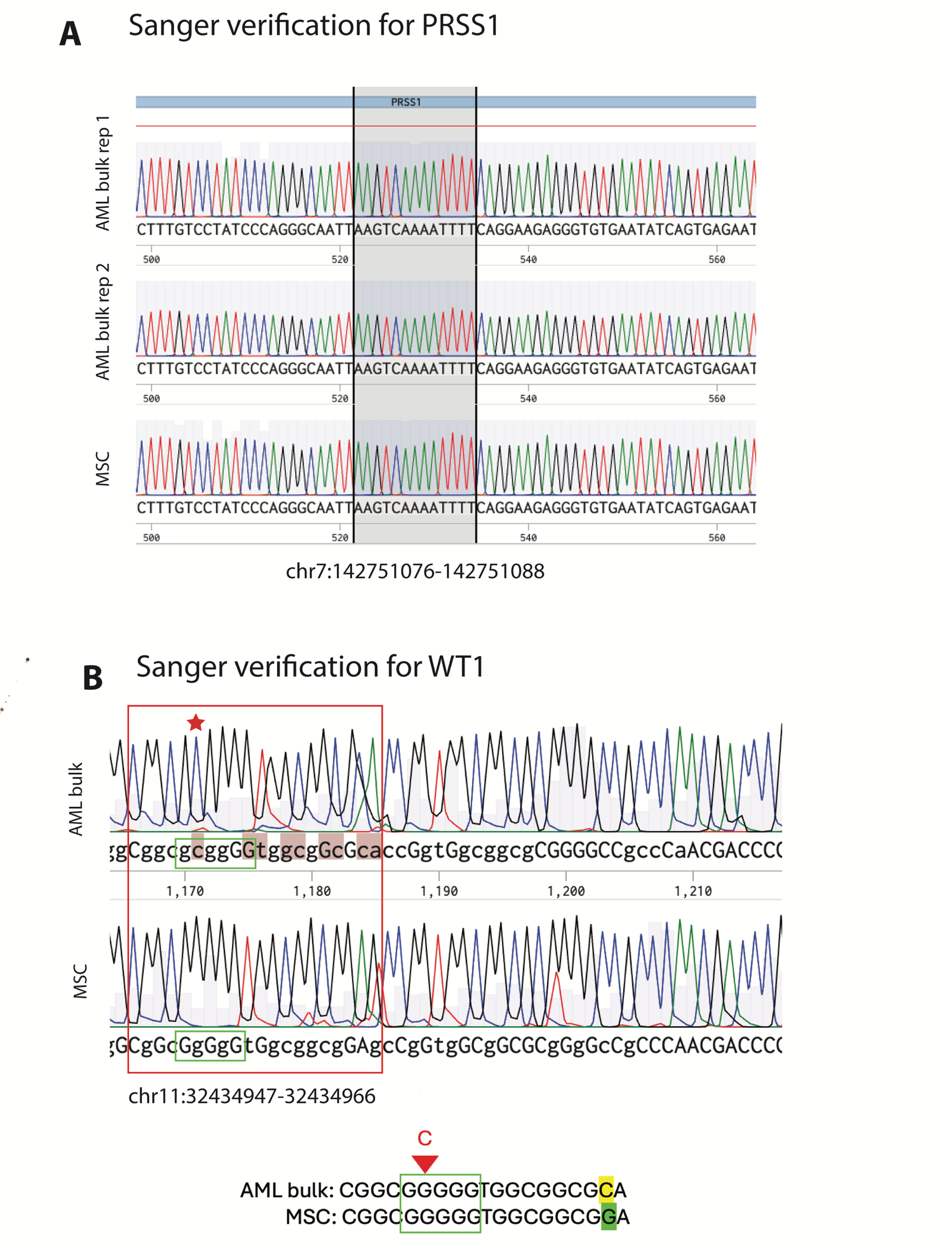

**Figure S15. Sanger sequencing results**. A – Sanger sequencing chromatogram for the region of interest of the PRSS1 gene (black frame), showing the absence of any mutations in AML bulk and MSC bulk samples. B - Sanger sequencing chromatogram for the region of interest of the WT1 gene (red frame); the place of the C insertion in the poly(G) sequence (green frame) is highlighted with the red star.
